# Spatial statistics for identifying and scoring immune clusters in high-plex profiles of primary prostate cancer

**DOI:** 10.1101/2025.09.21.677465

**Authors:** Ali Amiryousefi, Jeremiah Wala, Jia-Ren Lin, Brian William Labadie, Aishwarya Atmakuri, Zoltan Maliga, Eamon Toye, Kiranj Chaudagar, Madeleine S. Torcasso, Shannon Coy, Giuseppe Nicolo Fanelli, Brigette Kobs, Fabio Socciarelli, Andreanne Gagne, Eliezer M. Van Allen, Akash Patnaik, Peter Sorger

## Abstract

The spatial arrangement of immune cells in the tumor microenvironment (TME) varies widely, from dispersed to clustered and tumor excluded to infiltrating. Multiplexed spatial profiling is an effective means of characterizing tumor-infiltrating lymphocytes (TILs) and immune complexes such as tertiary lymphoid structures (TLS) in the TME. However, few approaches have been described for objectively parametrizing patterns of immune organization and assessing their association with biological or clinical variables. This makes it difficult to evaluate whether a set of tumors is relatively immunologically cold or hot. Here we describe an intuitive set of statistical tools (available in the R package, *tlsR*) for characterizing lymphocyte patterns in the TME of solid cancers. We apply *tlsR* to primary prostate cancer (PCa), which is often described as immunologically ‘cold’. Using a cohort of 29 radical prostatectomy specimens stratified into low Gleason-grade (LGG; n=15) and high Gleason-grades (HGG; n =14) we show that HGG PCa is significantly more infiltrated than LGG PCa with lymphocytes organized into B cell or T cell enriched immune clusters (BICs and TICs). A subset of these ICs have the B and T cell zonation and follicular dendritic cells characteristic of a bona fide TLS. HGGs are also enriched with ICs containing precursor exhausted T cells (T_pex_) and proliferating B cells and their tumor compartments harbor granzyme-B^+^ cytotoxic T cells in contact with cancer cells. Thus, far from being cold, a subset of HGG PCa has features associated with active immune surveillance, a finding with implications for emerging PCa immunotherapies.

## INTRODUCTION

The tumor immune microenvironment (TME) of solid cancers plays a critical role in tumor evolution and response to therapy(1–4). The TME is characterized both by its cellular composition and its spatial organization. Tertiary lymphoid structures (TLS)(5) represent one prominent spatial feature in which lymphoid-like structures (including germinal centers) form in non-lymphoid tissues in response to persistent immune stimulation. The presence of TLSs can be highly prognostic, most notably in melanoma(6) and colorectal cancer (CRC)(7). An extensive literature has therefore developed on the role of TLSs and other immune clusters (ICs) in cancer progression and immunotherapy, but a substantial limitation of these studies is that few objective criteria have been described for identifying an IC or TLS in peripheral tissues, and distinguishing ICs from more dispersed immune infiltration. Here we address this challenge by developing a suite of spatial statistical tools (available as an R package, *tlsR)* that objectively measure the absolute levels and relative distributions of B and T cells. We develop quantitative metrics for parameterizing ICs to identify significant associations between IC properties and tumor grade or other independent clinical variables and apply them to a tumor type in which the role of ICs, and the TME more generally, remains unclear: treatment-naïve prostate cancer (PCa).

Metastatic castrate-resistant prostate cancer (mCRPC) is often considered to be an immunologically “cold” tumor with a relative paucity of tumor-infiltrating immune cells and no established role for immunotherapy except in the rare case of microsatellite instability-high (MSI-H) tumors(8). In contrast, expert pathology review of H&E stained tissue sections nonetheless recognizes the presence of tumor-infiltrating immune cells in primary PCa from prostatectomies(9–11). Conventional IHC studies and more recently multiplexed IHC and multiplexed immunofluorescence (IF) have further confirmed the presence of tumor infiltrating lymphocytes(12–22), immunosuppressive regulatory T cells(23–25) and tumor-associated macrophages(26–28), among other cell types. Moreover, some PCas contain immune clusters ranging from small immune aggregates(29) to larger structures(30). For example, in a cohort of primary PCa selected for high PD-L1 expression, Calagua et al identified niches of MHC-II^+^ dendritic cells in spatial proximity to PD-1^+^ TCF1^+^ stem-like CD8^+^ T cells(29). These niches were associated with increased tumor-infiltrating lymphocytes (TILs), which is consistent with ICs as sites of antigen cross-presentation and tumor reactive T cells. Larger immune clusters have also been observed in both low- and high-grade PCa with multiplex IF, and at least a subset of these structures display canonical TLS features including proliferating CD20^+^ B cell follicles and follicular dendritic cells(30). These TLS-like structures are also identifiable in H&E images, with associated gene signatures supporting the identification of these primarily as T and B cell clusters(11). Moreover, recent multiplexed IF imaging of 13 primary PCa tumors identified cellular neighborhoods comprised of mast cells, regulatory T cells and M2-like macrophages, suggesting a spatial organization to immunosuppression programs previously identified by RNA-seq(31). Despite the growing body of evidence supporting demonstrating tumor-infiltrating immune cells in primary PCa, immunotherapeutic strategies remain largely unexplored in the localized PCa disease context.

These studies have limitations: (i) many involve single-channel IHC with limited molecular information (ii) others rely on tissue microarrays (TMAs) that lack power for spatial analysis(32) and (iii) in no case are criteria for identifying statistically significant aggregates of immune cells and patterns of T and B cell interaction well defined. We therefore combined *tlsR* and highly multiplexed cyclic immunofluorescence (CyCIF)(33) imaging to quantify the levels and relative distributions of immune cells in the TME of 29 radical prostatectomy specimens enriched for high Gleason grade (HGG; Gleason grade ≥ 4+4) tumors. These HGG tumors represent ∼20% of primary PCa tumors but half of PCa mortality(34,35). Applying tlsR to this cohort, we find that a substantial fraction of HGG PCa tumors have well organized ICs, some of which have the hallmarks of functional TLSs. Our findings reframe HGG PCa as a tumor with a structured and potentially actionable local immune response, while also providing a generalizable statistical framework for application across solid tumors.

## MATERIALS AND METHODS

### Imaging and data generation

The extended versions of marker names are provided in **Supplementary Table 1**, along with their associated antibodies and vendor information in **Supplementary Table 2**. All images were visually inspected by board certified pathologists, and segmentation mask dataset then obtained using MCMICRO(36), which generated the cell-feature table for each specimen. Areas of tissue with artifacts or poor quality(37) were eliminated from subsequent analysis. The specificity of staining was also checked visually, with emphasis placed on current morphology and signal to noise ratio (**Supplementary Fig. S1A**). Cell type identification was based on markers that had passed quality control (**Supplementary Fig. S1B**). We also used visual inspection to determine the marker intensity threshold above which a cell was called positive for that marker. For markers where good signal to noise was restricted to ICs (Ki67, TCF1, PD-1), we determined the positivity threshold only in these regions of interest. In all ICs in which gating on the above markers was possible, we observed a subset of Ki67^+^ B cells that also scored as TCF1^+^. While it is possible that these represent a recently identified self-renewing population of B-1 cells(38), we think it more likely that this is due to quality issues in the segmentation of immune cells in densely packed ICs, leading to signal spillover from neighboring cells(39). Thus, we judge the quantification of TCF1^+^ cell number to be reasonably reliable but the assignment to B cells a likely segmentation artifact.

### Approach to IC detection and parameterization

The methods we used to identify and characterize ICs are likely to be of general utility, and we therefore describe them here in greater detail. Two approaches were used to locate ICs: KNN clustering and DBSCAN; both have strengths. Starting with an immune cell centroid, a K nearest neighbor (KNN) algorithm classified cells based on the dominant property of K nearest neighbors (K=100 in the current study; this was not a sensitive parameter). KNN clustering was most effective with relatively dense and uniform clusters (typically when ≥50% of the 100 nearest cells had the same marker label). Visually, the distribution of T cells was more diffuse than that of B cells, and KNN clustering missed some of these aggregates. DBSCAN, an unsupervised method that groups together cells with similar marker labels in a manner that is robust to noise and outliers, was effective in this setting and detected 588 TICs, many of which were coincident or proximate to BICs, with significantly greater prevalence in HGG than LGG PCa. TICs contained ∼3-fold fewer cells than BICs and were generally less densely packed. BICs and TICs were found in association with each other in the most organized ICs, which had the properties of mature TLS, but TICs were also common independent of BICs. To our knowledge, this type of IC has not been extensively characterized outside of the setting of lymphomas and leukemias.

To quantify how TILs were dispersed throughout the tissue we applied a version of Ripley’s K-Function,(40) which is widely used in astronomy and more recently, in tissue imaging to detect deviations from homogeneous spatial distributions(41,42). The Centered L Function (CLF) is a variance-stabilized and centered version of Ripley’s K-Function that has the advantage of being density independent and homoscedastic. We used it to detect significant clustering or dispersion (regularity) of cells relative to complete spatial randomness (CSR). The CLF generates two readily interpretable whole-slide parameters, the clustering Intensity (CI) and the Effective Clustering Radius (ECR), which can be compared at a specimen or subgroup level for comparison against clinical variables (in this case LGG v HGG). The multitype Centered L-function (MCLF) is an extension that quantifies joint distributions of two cell types (T and B cells in this work). In PCa this demonstrated non-random T and B distributions (and co-occurrence) at radii as great as ∼1 mm, a value 2-4-fold greater than cluster radii obtained by KNN or DBSCAN.

To quantify the degree of organization of ICs within and across specimens we used independent component analysis(43), a classical dimensionality reduction method related to principal component analysis (PCA). ICA separates multivariate signals, the spatial distributions of cells in ICs in the current case, into statistically independent but not necessarily Gaussian components. ICAT, the normalized trace (the sum of the diagonal elements) of the ICA standard deviation matrix captures the degree of symmetry and the density of ICs in a single parameter. Direct comparison of ICAT values with expert pathology annotation of H&E images confirmed their correspondence (and thus, their biological interpretability) and showed that BICs and TICs were significantly more organized in HGG than LGG PCa. ICs with a high degree of organization (a low ICAT value) were more likely to exhibit features of germinal centers involved in B cell amplification and T cell programming including proliferating B cells, TCF1^+^ T cells, and CD21^+^ and CD23^+^ follicular dendritic cells. However, only a minority of these ICs had classical germinal center morphology in H&E images suggesting that this is too strict a criterion for ICs in which B and T cells have the potential to interact functionally(44).

### Handling of outliers and special samples

In the case of B cells, high variance in cell number was driven by a single outlier case (LSP12629), which had nearly 10-fold more B cells than the next highest tumor. This tumor exhibited an extremely dense CD20^+^ B cell infiltrate winding in and out of the tumor compartment. Upon image review, these cells showed specific membranous staining for CD20, abundant CD19 staining, and very little off-target marker expression (e.g., CD3d), confirming that they were likely to be B cells. The large number of B cells leads us to speculate that this patient had a concurrent or prior history of B cell lymphoma. To avoid bias in our analysis, we excluded this sample from the infiltration analysis. Additionally, two detected BICs in this patient, each exceeding 10^5^ cells in size, were excluded from all BIC-related analyses. Three slides from LGG cases (LSP12631, LSP12643, LSP12645) did not have a tumor compartment visible on the image, although their grade assignments were based on whole-block review by pathologists. These samples were also excluded from any analyses pertaining to the tumor, such as infiltration and proximity. In contrast, two LGG samples (LSP12603 and LSP12609) and one HGG sample (LSP12623) lacked B immune clusters; these were included in the overall analysis but excluded from the proximity chart.

### Comparison to colorectal cancer and infiltration

To compare the absolute magnitude of immune infiltration to another cancer type, we calculated the median CD8^+^ and CD20^+^ tumor immune infiltration from a cohort of 74 colorectal cancers (CRC) previously imaged and analyzed using equivalent multiplexed IF technology. This cohort encompassed 65 patients with mismatch-repair proficient (pMMR) tumors and 9 patients with mismatch-repair deficient (dMMR) tumors(45). Owing to deficiencies in nucleotide repair machinery, dMMR tumors have a significantly higher tumor mutational burden (TMB) and immune infiltrate compared with pMMR tumors and are primarily treated with first-line immunotherapy in the metastatic setting. The infiltration of B, CD4, and CD8 T cells was calculated for prostate and CRC samples by identifying all cells within areas annotated by expert pathologists, blinded to prior clinical pathology report or tumor-immune classification, as the tumor compartment and dividing by the total area in mm^2^ of that compartment.

### Statistical analysis

Multiple statistical tests were employed in this study, each selected based on the type of comparison and the most common assumptions about the data. Basic comparisons of central tendencies between two samples were performed using a *t*-test in cases with more than 25 data points per sample or when the normality assumption of the data distribution was not rejected by the Kolmogorov-Smirnov normality test; otherwise, the nonparametric Mann-Whitney U test was used. Multi-sample analyses were conducted with the nonparametric Kruskal-Wallis test. For assessing increasing trends in lymphocyte data, the Jonckheere-Terpstra trend test was applied. Infiltration values were compared with colorectal cancer counterpart values using the nonparametric Wilcoxon rank-sum test, while Levene’s test was used to compare variance between competing groups. The proportion z-test was used for comparing two proportion vectors, and for proportion vectors involving more than one subtype, dissimilarities were quantified using Jensen-Shannon divergence. For regression analyses, models were fitted with an intercept, and the overall goodness-of-fit was reported via the *F*-test (Fisher exact test) *P* value. In scenarios where the number of data points in one sample was more than twofold that of the other, the *R*^2^ value was also reported. The same handling of special samples (as discussed above) was applied to relevant regression analyses. In logarithmic regression analyses, samples with original zero values were excluded, as these were associated with suboptimal areas for gating specific markers. Box and violin plots, as well as regression fits, were generated based on Tukey’s outlier rule. UMAP projections were constructed using scaled marker intensities underlying the respective phenotypes, color-coded by the discrete assigned phenotypes, with parameters of 15 neighbors, a minimum distance of 0.08, and the Euclidean metric. The detailed mathematical derivation of ICAT, along with notes on the centered L-function (CLF), multitype L-function (MCLF), and derived summarization indices, are presented in **Supplementary Data File 1-2**. All packages used in this study, along with their respective applications and native environments, are provided in **Supplementary Table 3**.

### Detection of the BIC and TIC

As described in the documentation for *tlsR*, we used the K nearest neighbors (KNN) algorithm to detect B immune clusters (BICs). First, we identified clusters of cells where at least 50% of the 25-nearest neighbors were B cells. These were then expanded to include any neighboring immune cells (positive for CD11b, CD68, CD163, CD4, CD3d, CD8a, TCF1, FOXP3, PD-1, CD57, CD11c, GZB, CD15, HLA-DR, CD103, CD31, pTBK1, CD24, CD44, or CD206) if at least 20% of the 100 nearest neighbors of those immune cells were B cells. A minimum cluster eligibility cutoff of 300 cells was then applied to remove very small groups of cells. For T immune clusters (TICs), we used the DBSCAN algorithm, nucleated on T cells (positive for CD3d, CD4, and CD8), with a minimum cluster size cutoff of 100 cells and an expanding neighborhood radius of 50 μm to include proximate T cells.

### The spatial characterization with ICAT

To quantify the organization of all *n* cells with *x*, and *y* coordinates in each IC,

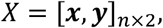

we employed independent component analysis (ICA), a computational method that decomposes input variables into the most statistically independent components(43);

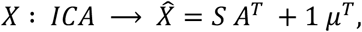

where *S* = [***s***_1_ ***s***_2_]_*n*×2_ is the independent components (transformed variables) and are uncorrelated and thus exhibit zero covariance in the off-diagonal elements of their covariance matrix 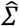. This is making the trace of the standard deviation matrix 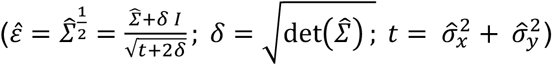 an indicator of spatial dispersion. Intuitively, when ICA is applied to the two-dimensional (*x, y*) spatial coordinates of cells, BICs with primarily circular morphology (e.g. sections from spherical TLSs) exhibit nearly perpendicular ICA component axes, in contrast to more oblique or acute axes for amorphous clusters. After normalization by the size of the cluster (*n*) and applying a constant multiplier (*α*), we used the trace of the standard deviation matrix of the resulting new coordinates:

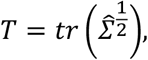

as a measure of the per-cell spatial standard deviation for each IC:

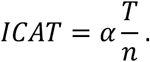

**Supplementary Data File 1** provides the expanded proof of this derivation and **Supplementary Table 4** lists the TLS identifiers, ICAT values, pathologist annotations, and relevant marker expressions.

### The K-function and co-clustering

The positions of cells in tissue can be modeled using a Poisson point process if their spatial locations are simplified to single (*x,y*) coordinates. Ripley’s K-function is used to quantify the spatial clustering or regularity tendencies of these points(46,47). Under complete spatial randomness (CSR), the K-function is expected to expand exponentially with increasing radius, whereas Besag’s L-function shows a linear relationship with radius(48). Consequently, subtracting this function by the radius (*r*) render this as a Centered L-function (CLF) which exhibits constant value under CSR. Upward deviations of CLF from this constant then indicate clustering intensity, while values below it suggest regularity(49). Formally, for distinct types of points *i* and *j* each with respective *n*_*i*_ and *n*_*j*_ size in a window *W*, and estimated isotropic edge-corrected K-function 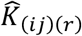, we obtained the variance-stabilized and centralized L-function estimate (shortly denoted ‘CLF’ here) as (see **Supplementary Data File 2**):

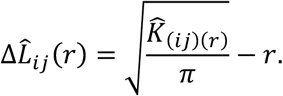

We further defined the positive envelope of this function by

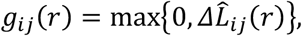

with largest radius above which no positive deviation persists as

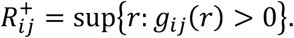

This allowed us first to characterize clustering intensity of the curve by

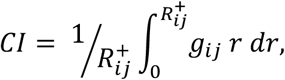

as the integral of the curve in its positive *r* domain. Furthermore, we derived R50 as the maximum radius at which the curve accounts for 50% of the positive integral area of the curve:

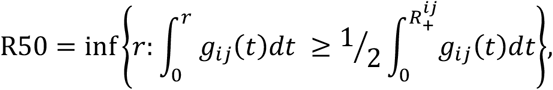

leading, second, to the effective clustering radius

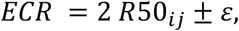

as the outer limit up to which significant clustering between points persist (2|*ε*| < binning size; **Supplementary Data File 2**). Together these two indices, captures the two properties of the curve underlying the meaningful clustering extent and its strength. For our cohort-based analysis, we computed the 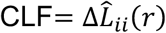 up to a 1.5 mm radius, using increments of 30 μm, separately for B and T cells. Tissue-wise curves were generated by averaging the estimated divergences of the CLF from CSR at each radius interval, across four bootstrap iterations of squared 9 mm^2^ tiles from all tissues. The bootstrapping step was included to mitigate bias from suboptimal tissue bracketing. Using identical parameters, the multitype L-function underlying the cases where *i* ≠ *j*, (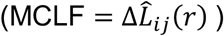) was estimated by counting the number of T cells within the radius of each B cell (**Supplementary Data File 2**). The 9 mm^2^ tile size was reverse-optimized by examining the behavior of the CLF and MCLF across a range of radii, ensuring capture of all interaction dynamics and a confident downward trend toward CSR within this window. For exemplary cases, we computed the CLF and MCLF over a 500 μm radius from each cell, using 10 μm increments. Analogously, co-clustering intensity (CCI) and effective co-clustering radius (ECCR) metrics were derived using the MCLF instead of the CLF (**Supplementary Data File 2**).

### Study approval

All studies involving human participants were conducted in accordance with the Declaration of Helsinki and applicable U.S. federal regulations. The collection and use of human tissue specimens and associated clinical data were approved by the Institutional Review Boards of the Dana-Farber Cancer Institute (IRB 01-045) and Northwestern University (IRB NU 00X3). Written informed consent was obtained from all participants prior to specimen collection.

### Data availability

All CyCIF images (across all channels), digitized H&E images, and cell segmentation masks are deposited to AWS servers at s3://lsp-public-data/amiryousefi-wala-prostate-2026/. The segmented data (feature tables), higher-level processed data, and additional resources required to reproduce the analyses are deposited in Zenodo at https://doi.org/10.5281/zenodo.20645947 and assigned a permanent DOI: https://doi.org/10.5281/zenodo.20645947.

### Code availability

The *tlsR* package containing the essential functions for detecting the BICs and TICs, as well as characterizing their organization with ICAT, and spatial point pattern analysis and plotting (M)CLF of the whole tissue, are deposited and accessible at CRAN at https://cran.r-project.org/web/packages/tlsR/ (DOI: 10.32614/CRAN.package.tlsR). The GitHub page for the tlsR package is at https://github.com/labsyspharm/tlsR, with complete functions definition and vignette walk-through at https://labsyspharm.github.io/tlsR/. All other code necessary to reproduce the results is released on GitHub https://github.com/labsyspharm/Amiryousefi-Wala-Prostate-2026 and are archived in Zenodo at https://doi.org/10.5281/zenodo.20645947.

## RESULTS

### Spatial immunophenotyping PCa reveals high lymphocytic infiltration in a subset of high-grade tumors

Gleason grade is the key diagnostic feature for risk-stratification and clinical decision making in treatment-naïve PCa. Gleason grade parameterizes tumor growth patterns, which range from well-formed and separated glands (Gleason 3) to a disorganized pattern of tumor cells as individual cells, cords, or solid sheets without gland formation (Gleason 5)(50). We obtained 29 PCas of varying Gleason grade from treatment-naïve radical prostatectomies performed at Northwestern Memorial Hospital between 2005 and 2014. Specimens were scored for grade, seminal vesicle involvement (SVI), perineural invasion (PNI), lymph node status, and tumor stage according to standard AJCC criteria (**Fig. 1A**). Initial spatial analysis was performed using the Gleason grade of individual tumor regions as the independent variable (1-13 subdomains per specimen and 99 total across 29 specimens); tumor region grade varied from Gleason 3+3 to Gleason 5. However, consistent with clinical practice, most analysis used the whole tumor score defined as the maximum Gleason grade across all histological sections examined. Whole tumor Gleason scores were dichotomized into “low Gleason grade” (LGG) corresponding to a maximal tumor Gleason grade of 3+4 and “high Gleason grade” (HGG) corresponding to Gleason 4+4 or any Gleason pattern 5. There were no patients with an overall Gleason grade of 4+3, although some tumor sub-regions exhibited Gleason 4+3 features. As expected, patients with high grade disease tended to have other high-risk clinical features, including lymph node involvement, SVI, and PNI, although these associations were not statistically significant.

**Figure 1.**
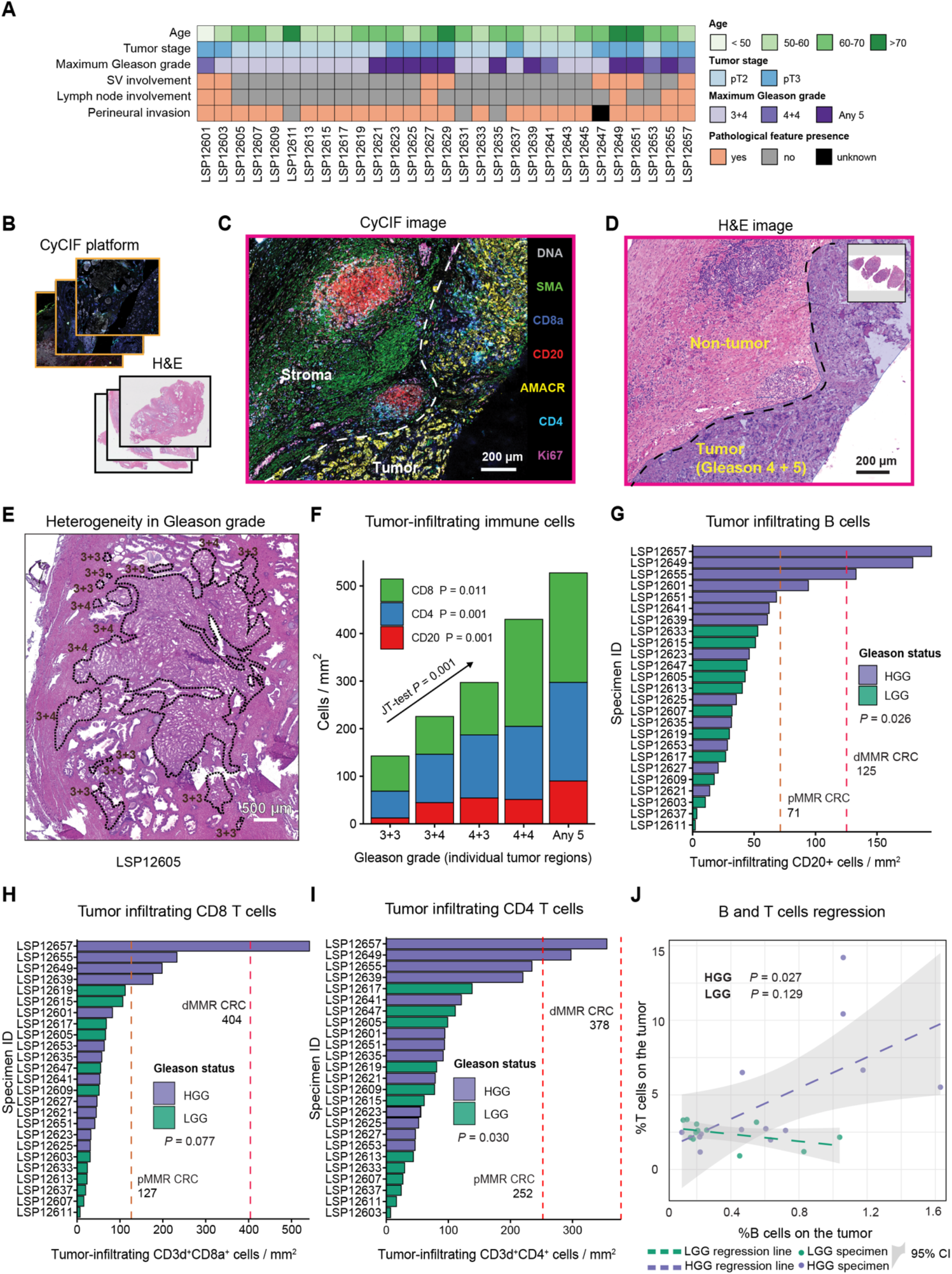
Whole slide CyCIF imaging reveals elevated B and T cell infiltration in high Gleason grade (HGG) PCa. **A**, Cohort overview with key clinical annotations. **B**, Experimental workflow: whole-slide cyclic multiplex immunofluorescence (CyCIF) and adjacent H&E imaging was performed on whole-slide radical-prostatectomy specimens. **C**, CyCIF image with selected markers shown; white dashed line separates the tumor and stroma compartment. **D**, Adjacent H&E image of the same specimen as in panel (**C**). **E**, Exemplar H&E image with individual tumor domains annotated with Gleason grades. **F**, Infiltration of B cells, CD4 T cells, and CD8 T cells per mm^2^ across Gleason grade groups G1–G5 (“3 + 3”, “3 + 4”, “4 + 3”, “4 + 4”, and “any 5”, respectively) within individually annotated tumor compartments (as in panel E) for all samples. The Jonckheere–Terpstra trend test (*P* value displayed) confirms an increasing lymphocytic immune-cell proportion with grade. Linear fit *P* values for each individual cell type are also shown (*F*-test). **G**, Density of intratumoral CD20^+^ B cells (cells per mm^2^) by Gleason grade. The one-sided Mann-Whitney U test *P* value compares high-vs. low-grade disease; dashed lines indicate median reference densities in colorectal cancer as indicated. Point-wise one-sided Wilcoxon test *P* values for adjacent grade comparisons are indicated. **H**, Similar to (**G**), density of intratumoral CD8^+^ T cells (cells per mm^2^) by Gleason grade. **I**, Similar to (**G**), density of intratumoral CD4^+^ T cells (cells per mm^2^) by Gleason grade. **J**, Regression fit and confidence intervals of the proportion of the intratumoral T cells (CD3d^+^) versus B cells (CD20^+^) stratified by Gleason status. *P* values represent the *F*-test of the fit.

21-marker CyCIF(51) spatial profiling was performed on whole slides (∼10^6^ cells per specimen) rather than TMAs, providing the spatial power needed to characterize large immune clusters (IC)(32) (**Fig. 1B-C; Supplementary Fig. S1A; Supplementary Table 1**). Images were processed using routines in MCMICRO(36) to generate feature tables comprising the x-y coordinates of each segmented cell (2.7 × 10^7^ in total) and the mean intensities of each marker within either a nuclear or whole-cell segmentation mask. Per-cell marker intensities were gated to generate positive and negative classes (see **Methods**) and cell types were then identified based on binarized marker combinations (**Supplementary Fig. S1B**). To distinguish the tumor compartment and surrounding non-cancerous tissue, tumor boundaries were identified using H&E images by board-certified pathologists who were blinded to the clinical pathology report or tumor-immune classification status; most specimens were found to contain multiple tumor domains (1 to 13) separated by non-tumor prostate tissue ranging in size (in a 2D section) from as few as 44 cells in a tumor area of ∼0.0078 mm^2^ to 2.3 × 10^6^ cells in an area of 320 mm^2^ (**Fig. 1D-E; Supplementary Fig. S1C**); this corresponded to ∼1 to 95% of the specimen area. Uniform manifold approximation and projection (UMAP) visualization on cell marker intensities (**Supplementary Fig. S1D**) confirmed the expected discrimination of immune, stromal and tumor cell types with specimen-level differences confined primarily to the tumor compartment.

When we quantified the proportions of segmented cells within pathologist-defined tumor domains expressing the T cell markers CD4 or CD8 or B cell marker CD20, T cells were found to be ∼8-fold more abundant than B cells. When individual tumor regions were stratified by Gleason grade group, a monotonic increase was observed in all three immune cell populations with increasing tumor grade (Jonckheere-Terpstra (JT) test, *P* = 0.001; **Fig. 1F**). The mean density of tumor-infiltrating CD20^+^ (B) cells varied almost 90-fold in the PCa cohort (∼2 cells/mm^2^ to ∼194 cells/mm^2^; **Fig. 1G**) and was significantly higher in HGG than LGG (Mann-Whitney U test, *P* = 0.026). As a reference, we compared immune cell densities in PCa to those in dMMR colorectal cancer (CRC)(45), a classically immunologically “hot” tumor.(52) In HGG PCa, B cell density approached that of dMMR CRC, a B cell rich tumor in which TLS abundance is predictive of slow tumor progression and ICI response. The density of CD3d^+^ CD8a^+^ cells (hereafter CD8 T cells) varied 75-fold across the PCa cohort (∼7 to ∼542 cells/mm^2^; **Fig. 1H**) but was significantly greater in HGG (mean = 123 cells/mm^2^) than in LGG PCa (mean = 48 cell/mm^2^); it was also significantly lower than either pMMR (204 cells/mm^2^) or dMMR (477 cells/mm^2^) CRC. While CD3d^+^ CD4^+^ cells (CD4 T cells) were significantly more prevalent in HGG than LGG PCa (Mann-Whitney U test, *P* = 0.030) they were also ∼3-fold less dense than in CRC (**Fig. 1I**). The proportion of all T cells in HGG PCa tumor compartments was nonetheless positively correlated with the proportion of tumor-infiltrating B cells (*F*-test, *P* = 0.027) whereas there was no correlation in LGG tumors (*F*-test, *P* = 0.129; **Fig. 1J)**. Similar results were obtained when relationships between B cells and either CD4 T or CD8 T cells were evaluated individually (**Supplementary Fig. S2A**). These results demonstrate that HGG PCa is infiltrated by B cells at a density approaching that of immunologically “hot” dMMR CRCs but has fewer T cells. Correlation in the levels of T and B cells in HGG PCa is consistent with formation of organized ICs in which T and B cells interact.

### Immune clusters in high-grade PCa are more numerous and localized to the tumor region

Visual inspection of CyCIF images suggested that some B cells in HGG PCa were organized into compact ICs (**Fig. 2A, Supplementary Fig. S2B)** but not all B cell clusters had classical germinal center morphology as judged from adjacent H&E images. To identify BICs in an objective manner we used a K nearest neighbor (KNN) classifier with a minimum cluster size set empirically at 300 cells (see **Methods**). This yielded 257 BICs across 29 specimens (note that the term “BIC” is intended to convey non-random spatial patterning of cells rather than any specific biological function; we expect functional TLSs to be both BICs and TICs). Some BICs had a compact round or ellipsoidal shape whereas others were less dense and more irregular; some BICs were located within the tumor domain and others lay in adjacent tissue (**Fig. 2B; Methods**).

**Figure 2.**
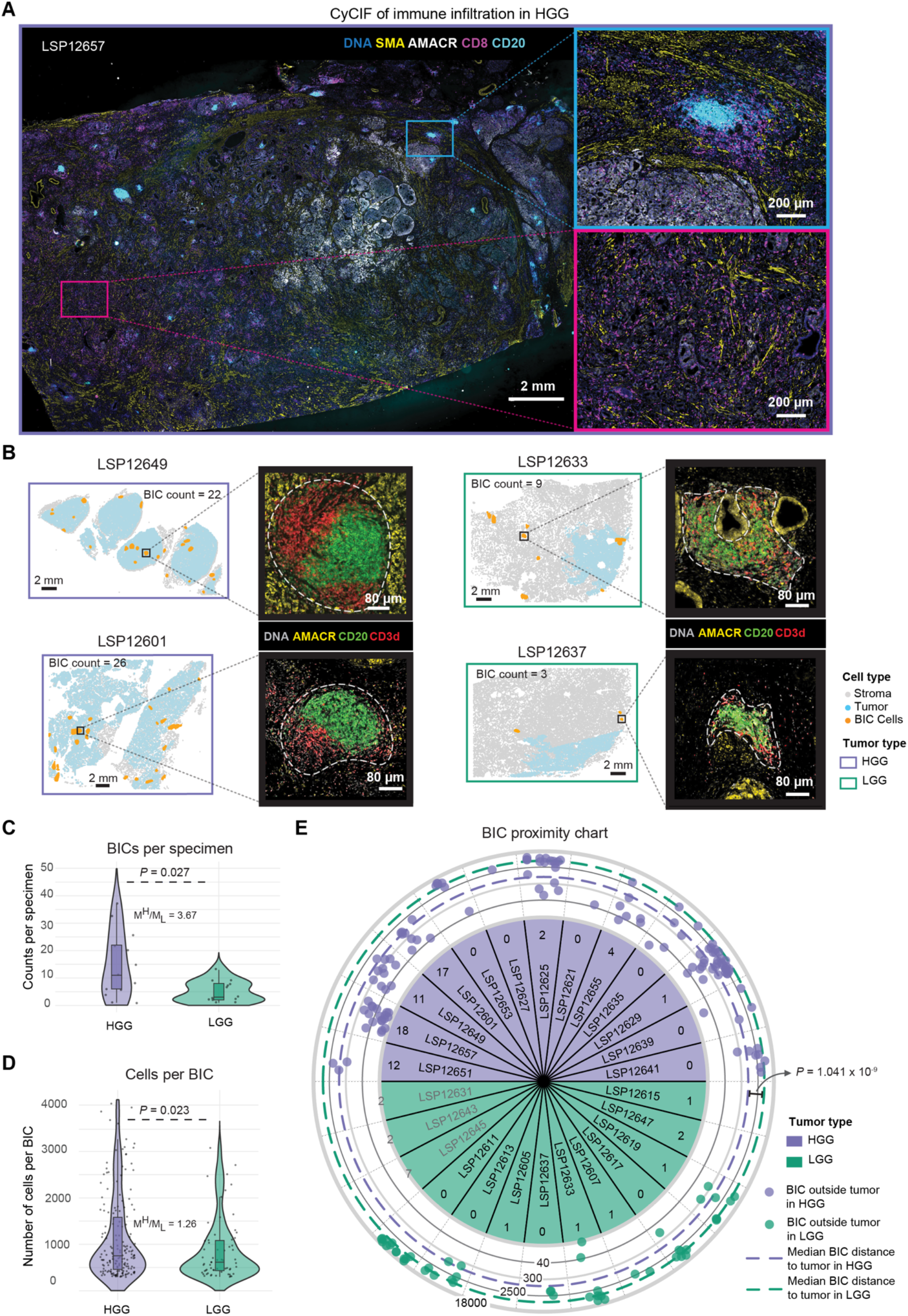
B cell immune clusters (BICs) are larger, more numerous and more tumor-proximal in high-grade PCa. **A**, Whole-slide CyCIF image of an exemplar HGG case with high immune infiltration, displaying selected markers for nuclear DNA, smooth-muscle actin (SMA), CD20, and CD8. A region with a BIC and a region with dense CD8^+^ T cell infiltration is shown in light blue and purple windows, respectively. **B**, Exemplar pseudo-images (cells represented as circles) for two HGG (left) and two LGG (right) specimens depicting non-tumor cells (gray), pathologist-defined tumor region (light blue) and detected BICs (orange). CyCIF images with AMACR, CD20, and CD3d markers for selected BICs. The approximate boundary of each BIC is denoted by the white dashed line. **C**, Number of the detected BICs per sample stratified by tumor grade. *P* values are from Mann-Whitney U test. The ratios of median values comparing HGG and LGG (M^H^/M_L_) are also shown. **D**, Cell counts per BIC size stratified by tumor grade with similar test and comparison to (**C**). **E**, Schematic depiction of the logarithmic distance from each BIC to the tumor compartment for extra-tumoral BICs (outer rings in μm), and the total count of intra-tumoral BICs for intratumoral BICs (inner ring). *P* value represents the Mann-Whitney U test of the BIC-tumor distance comparing HGG and LGG PCa. Three samples without annotated tumor regions on our slide are highlighted in grey.

We found that BICs in HGG were ∼4-fold more abundant (median BICs per sample of 11.5 for HGG and 2.5 for LGG, Mann-Whitney U test, *P* = 0.027; **Fig. 2C**) and slightly larger than in LGG PCa (median cells per BIC of 750 for HGG and 610 for LGG, Mann-Whitney U test, *P* = 0.023, **Fig. 2D; Supplementary Fig. S2C**). In HGG PCa, BICs were twice as likely to reside within the tumor compartment as in LGG (Mann-Whitney U test, *P* = 0.019). BICs outside of the tumor compartment were ∼7-fold closer to the tumor boundary in HGG than LGG (median tumor to BIC distance = 799 μm in HGG and 5366 μm in LGG; Mann-Whitney U test, *P* = 1.04 × 10^-9^; **Fig. 2E; Supplementary Fig. S2D**). However, this comparison is likely confounded by the relative sparsity of non-tumor tissue in HGG specimens relative to LGG specimens (that is, the lower number of “distant” BICs in HGG specimens may arise simply because the relevant tissue was not present in the samples). Overall, we conclude that, in HGG PCa, BICs were both more frequent and more likely to be spatially associated with the tumor compartment than in LGG tumors.

The impact of tumor infiltrating B cells (TIL-Bs) on tumor progression is complex, with reports of both beneficial effects due to antibody production or antigen presentation and adverse effects, due to cytokine-mediated TME reprogramming and suppression of T cell responses.(53) Unfortunately, our cohort was not powered to examine the association between BICs and clinical outcomes: with 14 HGG cases, an association between BICs and outcome would be detectable only if the data show near-perfect separation. We nonetheless asked whether the presence of high numbers of BICs was associated with a need for treatment post-surgery, excluding adjuvant radiation. At a median follow-up of 147 months, six patients (all with HGG) required additional treatment for prostate cancer. Among these six, three had a BIC count greater than the mean for HGG tumors; thus, BIC status was not significantly associated with receipt of additional treatment (p = 0.245, Fisher’s exact test). Fortunately, recent work from our groups has demonstrated that the presence of ICs at the time of radical prostatectomy, as detected by use of machine learning classifiers on H&E images, is associated with improved metastasis-free survival exclusively in patients with high-grade disease(54).

### Immune clusters in high-grade prostate cancer are organized and proliferative

To quantify differences in BIC morphology and density (**Fig. 3A**) we developed an approach based on independent component analysis (ICA)(43); this maps the *x*, y positions of *n* specific cell types (in the current case, cells in the BIC) onto maximally independent axes generating new 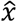, ŷ positions such that the linear correlation between these axes tends to zero; as a result the covariance matrix 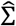 is nearly diagonal (see **Methods**). The normalized trace (*tr*) of the standard deviation matrix of the independent components,

**Figure 3.**
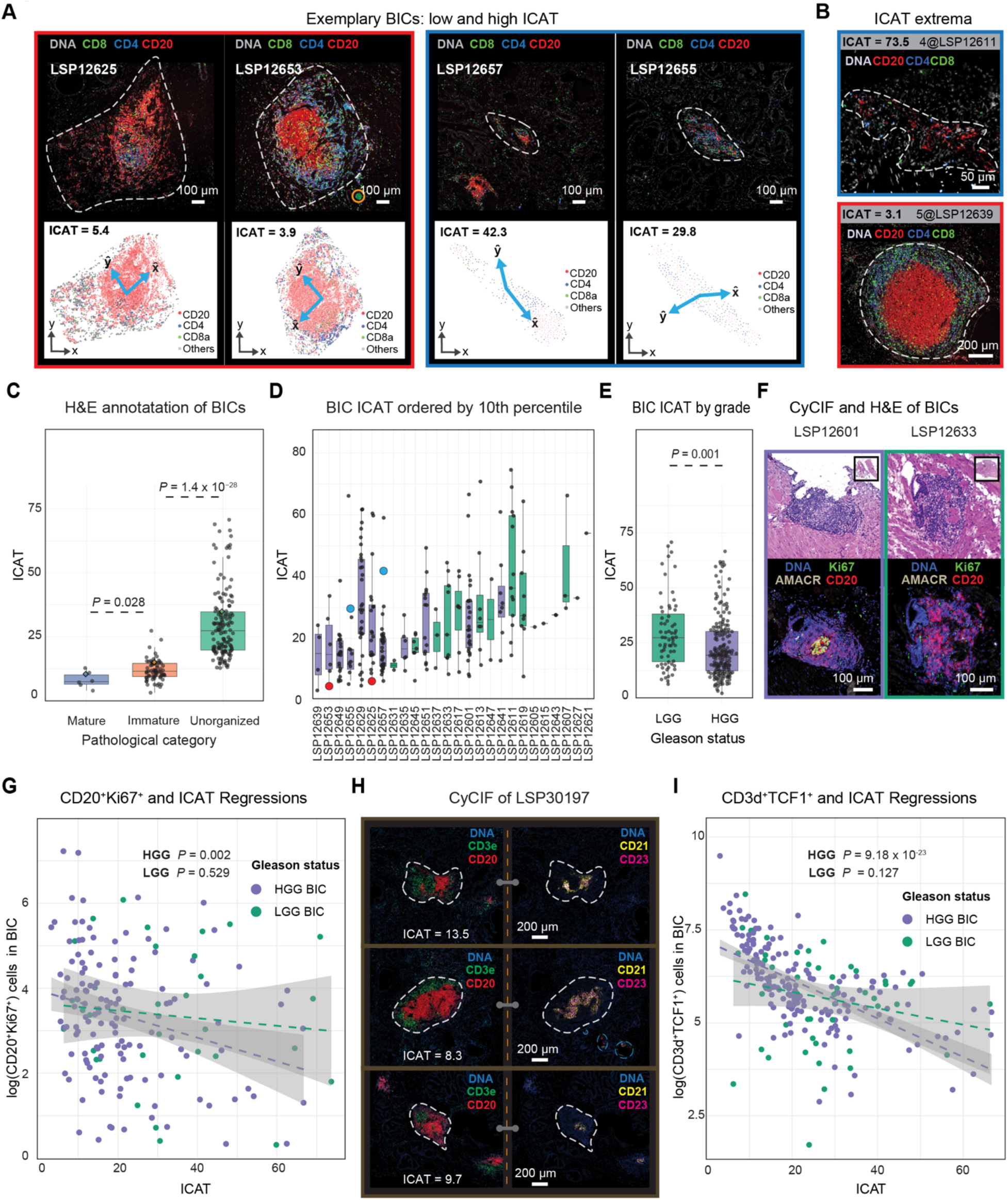
B cell clusters in high-grade PCa are spatially organized and harbor TLS-like immune-cells. **A**, CyCIF (top; approximate BIC boundary represented by white dashed line) and pseudo-images (bottom; cells are circles) of two highly organized (left panel) and disorganized (right panel) BICs. The blue arrows represent the independent components as determined by ICA analysis. Image artefacts are circled in orange. **B**, CyCIF images of the BICs with the highest (top; least organized) and lowest ICAT values (bottom; most organized). **C**, BIC ICAT scores stratified by pathologist classification. *P* values represent Mann-Whitney U tests. **D**, Distribution of ICAT values per sample, ordered by 10^th^ percentile of ICAT score. The four BICs in (**A**) are marked with red and blue for the left and right panel, respectively. **E**, Distribution of ICAT values by high-grade (HGG) versus low-grade (LGG), shown with *t*-test *P* value. **F**, H&E (top) and CyCIF (bottom) of exemplar BICs with (left) and without (right) a Ki67^+^ proliferative germinal center. **G**, Regression fit and 95% confidence interval of proliferative B cells (CD20^+^ Ki67^+^) compared with ICAT values per BIC with *R*^2^_HGG_ = 0.07 and *R*^2^_LGG_ = 0.01 and the *F*-test *P* value for regression fits. **H**, CyCIF images of three TLSs as characterized with follicular dendritic markers CD23 and CD21, depicted with the corresponding ICAT values for an external sample (LSP30197). Image artifacts are encircled with dashed light blue lines. **I**, Regression fit and 95% confidence interval of TCF1^+^ T cells (CD3d^+^ TCF1^+^) compared with ICAT values per BIC with *R*^2^_HGG_ = 0.49 and *R*^2^_LGG_ = 0.05 and the *F*-test *P* value for regression fits.

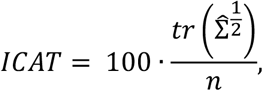

then quantifies spatial dispersion. Low ICAT values are associated with dense and symmetric BICs, and high values with BICs having fewer immune cells and arranged in irregular shapes (see methods; **Fig. 3A-B, Supplementary Data S1**). We validated the ICAT metric via expert pathologist review of adjacent H&E images. Among 257 BICs, six (2.2%) were judged from H&E images to have a high likelihood of representing a mature TLS with a germinal center; all were intra-tumoral BICs from HGG cases. An additional 72 (27%) of BICs were classified as immature TLSs, and 65/72 of these (90%) were found in HGG tumors. The remaining 181 BICs were classified as disorganized immune clusters with atypical and sparse cellular aggregates (**Supplementary Fig. S3A**). ICAT values recapitulated manual classification: mature BICs corresponded to ICAT= 8.1 ± 3.2, immature BICs to ICAT = 11.9 ± 4, and disorganized BICs to ICAT = 30 ± 14 (Kruskal-Wallis test of similarity, *P* = 2.2 × 10^-16^; **Fig. 3C**). As judged visually from H&E images, mature or immature TLSs were ∼4-fold more prevalent in HGG than in LGG PCa and BICs from HGG PCa were associated with significantly lower ICAT values than BICs from LGG tumors (Mann-Whitney U test, *P* = 0.001; **Fig. 3D-E**). From an analytical perspective, we conclude that ICAT values correspond well to histopathology classes without having been trained on them; ICAT is also an integrated parameter of spatial organization that can easily be correlated with continuous or categorical biological properties (e.g. LGG v HGG). More complex machine learning methods we have used previously(55), such as spatial LDA(56) do not have this property. From a disease perspective, we conclude that BICs in HGG PCa are significantly more organized than those in LGG PCa.

TLS represent sites in which B cells can undergo selection, amplification, affinity maturation, and eventual production of antibodies against tumor antigens.(44) A common measure of this process is the presence of proliferating Ki67^+^ B cells(57). Across all BICs, we found that ∼3% of B cells were Ki67^+^. However, BICs in HGG exhibited a significantly increased Ki67^+^ proportion as compared to LGG BICs (**Fig. 3F, Supplementary Fig. S3B**). A significant negative relationship was also observed between the log number of Ki67^+^ B cells and ICAT values in HGG tumors themselves (*F*-test, *P* = 0.002 and *P* = 0.529 for HGG and LGG, respectively; **Fig. 3G**). Moreover, detailed CyCIF characterization using an expanded TLS-focused panel (**Supplementary Table 1**) of an HGG PCa having ICs with ICAT values in the lower quartile (< 13.7 = Q_1_, corresponding to mature and immature histological classes; **Fig. 3H**) showed that organized BICs contained foci of CD21^+^ and CD23^+^ follicular dendritic cells (FDCs). The presence of FDCs is a key feature of functional germinal centers. We conclude that low ICAT ICs are more likely to have proliferating B cells and FDC foci, two characteristics of functional TLSs.

TLSs also promote programming and amplification of “stem-like” T cells, which can be identified based on expression of the TCF1/TCF7 transcription factor(58). BICs in our PCa cohort contained ∼1.3-fold as many T as B cells and 48% of these T cells were TCF1^+^ (among these, 8% were also Ki67^+^, **Supplementary Fig. S3C**). The log-number of TCF1^+^ T cells was negatively correlated with *ICAT* values for the HGG group (*F*-test, *P* = 9.18 × 10^-23^ for HGG and *P* = 0.127 for LGG; **Fig. 3I)**. The number of Ki67^+^ B cells and TCF1^+^ T cells was positively correlated in log-log space (*F*-test, *P* = 1.29 × 10^-8^ for HGG and *P* = 0.033 for LGG). Another way of viewing this result is that Ki67^+^ B and TCF1^+^ T cells were highly enriched in BICs having ICAT values less than its 1^st^ quartile (Q_1_ = 13.7) **(Supplementary Fig. S3D-E)**, and thus, segregation on ICAT interquartile value alone (<Q_1_ = 13.7 and >Q_3_ = 32.4) was effective at discriminating BICs with proliferating B cells and stem-like T cells in both HGG and LGG PCa. (**Supplementary Fig. S3F**). These data show that BICs in HGG are not only more numerous than in LGG PCa, but also significantly more organized (having lower ICAT values) and that organization is correlated with greater prevalence of proliferating Ki67^+^ B cells and TCF1^+^ stem-like T cells. These are features of a dynamic and active immune response involving both B and T cells.

### High-grade prostate cancers exhibit dense T cell clusters closely associated with B cell aggregates

The analysis of ICs described above focused on B cells because these are key constituents of germinal centers; however, inspection of images also revealed the presence of ICs primarily or exclusively containing T cells. As a first step in evaluating the extent of B and T cell co-clustering, we generated CyCIF contour density maps for B, CD4, and CD8 T cells (**Fig. 4A**). This revealed the presence of T cell-rich immune clusters lacking B cells (TICs; **Fig. 4A** panel b) as well as B cell specific BICs and ICs in which B and T cells were coincident (as described above). Examination of TICs in adjacent H&E images showed that they corresponded to lymphocytic aggregates that were indistinguishable by human inspection from less organized BICs (**Fig. 4B, Supplementary Fig. S3A**). To enumerate and characterize TICs, we performed DBSCAN on CD4 and CD8 T cells (**Methods**)(59). DBSCAN is a non-parametric density-based clustering algorithm that groups points (cells in this case) that meet a specific density per unit area criteria. It is effective at locating clusters with varying geometries and lack of clear centrality, which was typical of TICs in PCa (**Fig. 4B & D;** panel a).

**Figure 4.**
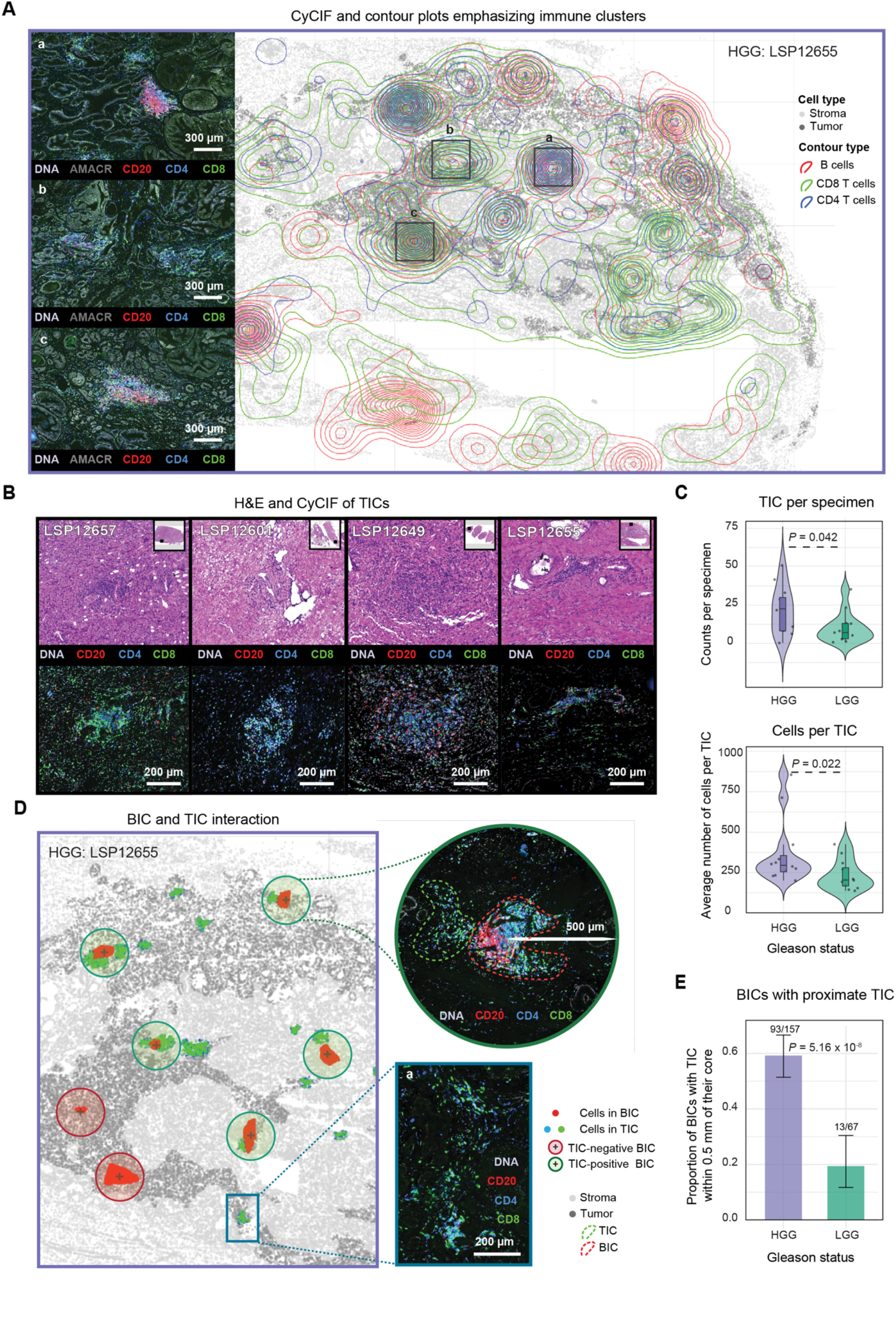
T cell clusters (TICs) are abundant and spatially co-localized with B cell clusters in high-grade PCa. **A**, Contour plot of an HGG specimen with high CD20^+^ and CD4^+^-CD8^+^ density areas (marked as *a* and *c*) and dense areas of CD4^+^ and CD8^+^ without apparent CD20^+^ intensity (marked as *b*). **B**, Examples of four post-CyCIF H&E images (top) and CyCIF images (bottom) denoting the T cell-clusters (TICs). **C**, Mean cell counts (top) and number (bottom) of TICs per sample, stratified by tumor grade (*P* values represent Mann-Whitney U test). **D**, Pseudo-image (cells are dots) of a HGG specimen depicted with BICs (red) and TICs (green), overlaid with green and red circled area denoting BICs with (green) or without (red) TIC within 500 μm from BIC core. Shown are exemplar corresponding CyCIF images with BIC (red dotted line) and TIC (green dotted line) circled (top) and a TIC without a corresponding BIC and particular organized shape (panel *a* bottom). **E**, Bar plot and 95% confidence interval of the proportion of BICs proximal a TIC (500 μm), stratified by tumor grade (*P* value denotes proportion test).

We identified 588 TICs across the PCa cohort, and these were on average ∼3-fold smaller (median 213 cells per TIC v. 713 cells per BIC; Mann-Whitney U test, *P* = 0.026) and 4 times “less organized” than BICs as measured by ICAT value (median TIC ICAT = 94; median BIC ICAT = 22, Mann-Whitney U test, *P* = 2.2 × 10^-16^). Although smaller, TICs were two times more numerous than BICs (588 v 257) and were significantly more abundant and larger in HGG as compared to LGG tumors (median of 7 TICs per specimen of 174 cells each in LGG vs. 24 TICs per specimen of 272 cells each in HGG; Mann-Whitney U test, *P* = 0.042 and *P* = 0.022, respectively; **Fig. 4C, Fig. 2C**). TICs were more likely to lie within or immediately adjacent to the tumor domain (HGG vs. LGG, Mann-Whitney U test, *P* = 3.26 × 10^-12^; **Supplementary Fig. S4A-C**). Thus, HGG PCa has larger and more organized TICs than LGG PCa, some of which are spatially associated with BICs. The study of ICs in tissue has focused primarily on structures containing both BICs and TICs – which appear to correspond to TLS – but our results show that T cell only TICs represent another common assembly of immune cells in association with PCa grade; further analysis of TIC properties and biological significance is therefore warranted.

Interaction of T and B cells is necessary for antigen presentation by B cells to helper T cells, co-stimulatory signaling, cytokine exchange, and the formation of immune synapses that facilitate B cell maturation and T cell activation(60–62). Thus, spatial proximity of B and T cell clusters is likely to be reflective of an active immune response. In PCa we observed that the numbers of BICs and TICs per specimen were positively correlated in HGG (*F*-test, *P* = 0.04) but not LGG (*F*-test, *P* = 0.63; **Supplementary Fig. S4D**). In a mature TLS, B and T cells are not fully intermixed, but instead exhibit “zonation”; this typically manifests itself as a T cell crescent surrounding a B cell core(63). We observed that the proportion of BICs with a TIC within 500 μm of the BIC center was 3-fold higher in HGG than LGG tumors (∼60% vs. ∼20%; two-proportion z-test, *P* = 5.6 × 10^-8^; **Fig. 4D-E**). Thus, TICs in HGG are more abundant, larger in size, and more likely to be associated with BICs; however, TICs not proximate to BICs are also found in PCa.

### B and T cell spatial organization in prostate cancer extends beyond discrete clusters

To better understand the organization of lymphocytes independent of their inclusion in immune clusters, we developed a complementary approach to quantifying the spatial organization of cells; this employed free-range spatial point pattern analysis across whole-slide images. For cell type *i* and *j* (e.g. B and T cells), and radius *r*, the multitype Centered L-function 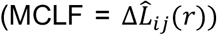 is the variance-stabilized and centered version of the more familiar edge-corrected Ripley’s K, 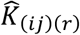. By computing the degree of divergence from complete spatial randomness (CSR) over varying radii, the Centered L-function 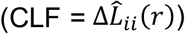 characterizes the degree of spatial interaction in sets of objects (different cell types in this case, **Methods; Supplementary Data S2, Fig. 5A**). Divergence is either positive (upward from the X axis, with CLF= 0 corresponding to CSR), representing greater clustering (red region in **Fig. 5A**) or negative, representing regularity (blue region); regularity refers to a pattern in which cells are more dispersed than would be expected by random assortment (**Methods**). We parameterized the CLF by (i) clustering intensity (CI) which is the averaged integral of the positive values of the function and an (ii) effective clustering radius ECR which is the point at which the CLF reaches zero and complete spatial randomness is reached. To make this estimate robust, the ECR is defined as the point at which the CLR crosses the upper ninety-five percent confidence interval for the estimate of CLF = 0 **(Fig. 5A, Supplementary Fig. S5A-D, Methods, Supplementary Data S2**). In contrast to the clustering methods described earlier, the CLF does not identify individual clusters but instead is ideal for quantifying spatial distributions irrespective of density. The ECR then provides an estimate of the radius (*r*) up to which a significant deviation from randomness is observed.

**Figure 5.**
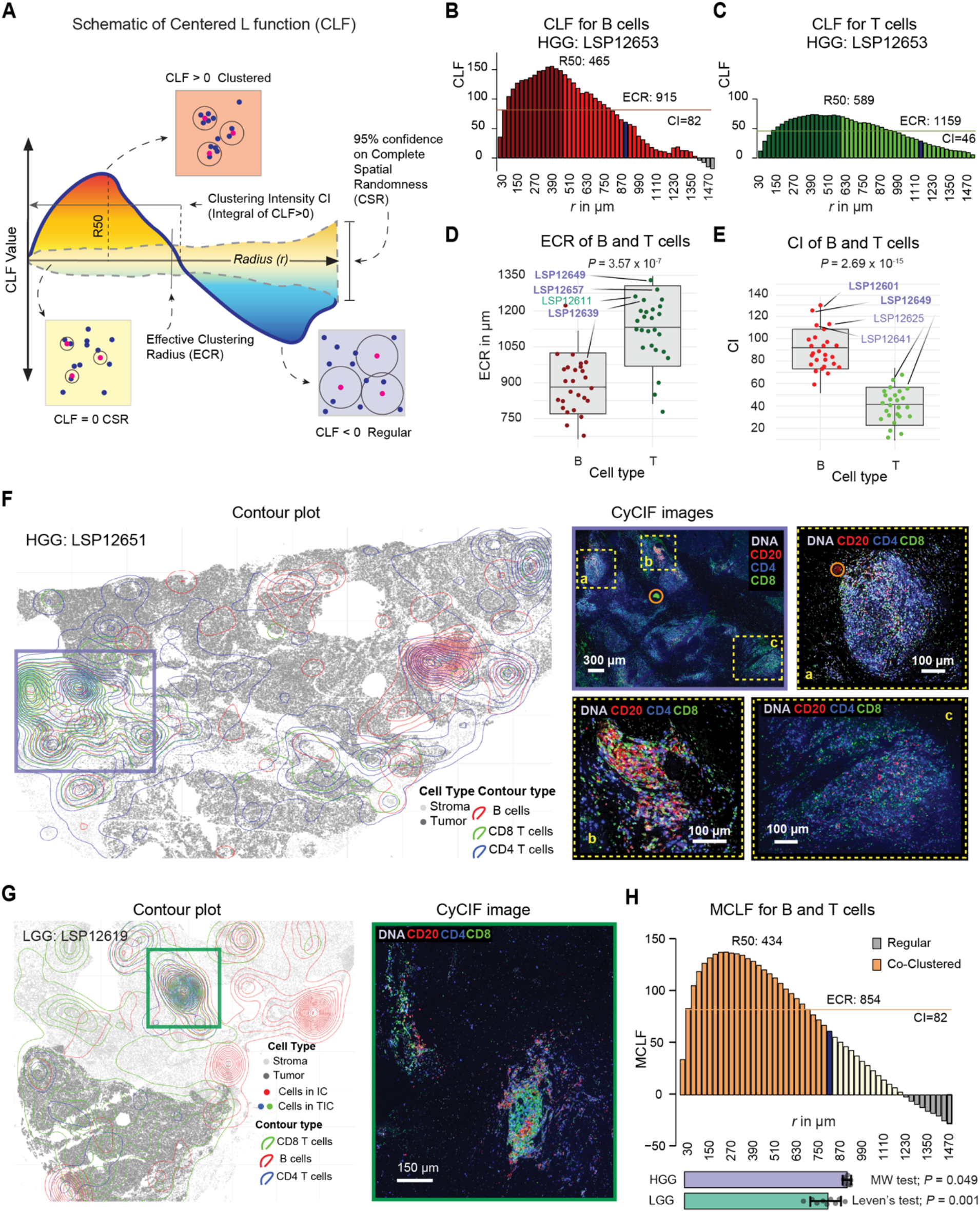
HGG exhibit increased B and T cells interaction on larger length scale. **A**, Schematic depiction of the centered L-function (CLF) over a span of radii (*r*), showing the complete spatial randomness (CSR) line and the expanding confidence interval around it. As a function of *r*, the example CLF begins with mild clustering at small radii, peaks at extreme clustering near *R*50 radius, crosses the CSR line into regularity in the CSR region, and then at higher radii exhibits non-random anti-clustering (regularity), returning to the CSR confidence area; R50: the maximum radius below which 50% of the positive integral accumulates, effective clustering radius (ECR): the maximum radius where significant positive clustering exists. **B**, The averaged CLF for B cells in 9 mm^2^ areas on the tissue, using fourfold systematic bootstrapping, showing peak clustering at radius of 455 μm. **C** Similar CLF and indices as in (**B**), but for T cells. **D**, ECR between T and B cells across the cohort (*P* value represents Mann Whitney U test). The four highest values in each boxplot are annotated with specimen identifiers for HGG (purple) and LGG (green), with specimens in the top 10th percentile for infiltrating CD20^+^ or CD8^+^ cells in the tumor across the cohort shown in bold. **E**, Clustering intensity (CI) between T and B cells across the cohort, with similar annotations as in (**D**) but *t*-test. **F**, Contour plot of an exemplar HGG sample highlighting large B and T cell co-clustering areas, with an overview of the CyCIF image area and zoomed-in sub-areas colored by CD20, CD4, and CD8 markers. Image artifacts are circled in orange. **G**, Contour plot of an LGG sample with an enlarged CyCIF image area showing a dense T cell community. **H**, The integrated multitype CLF (MCLF) as an average co-clustering survey of the whole cohort on 9 mm^2^ areas, depicting the interaction function for co-clustering (or co-regularity) of B and T cells similarly to (**B**), with overlaid co-clustering intensity (CCI) as the integral of the positive area of the MCLF and effective co-clustering radius (ECCR) as the maximum radius where significant positive co-clustering between B and T cells exists. The horizontal bar plot at the bottom shows the average ECCR for HGG and LGG samples with respective 95% confidence intervals, with differences in means and variances quantified by *P* values from the Mann-Whitney U test and Levene’s test, respectively.

With the CLF in hand, we asked whether B and T organization extends beyond the length scale of immune clusters. We computed the CLF using a 9 mm^2^ sliding square window based on exploratory studies that showed it to be ∼3 times greater than the largest ECR (**Supplementary Fig. S5E, Methods**). The CLF was summed across these windows to generate a per-tissue parameterization of immune cell organization; functions were also averaged across the cohort to generate summary statistics (**Methods**). By way of example, for HGG specimen LSP12653 (**Fig. 5B-C**), the B cell CLF rose to 150 at a radius of ∼455 μm and then fell monotonically to CSR at ∼1.1 mm, yielding a CI of 82 and ECR for B cells of 915 μm (**Methods, Supplementary Data S2**). For T cells, the CLF was similar in shape, yielding a CI of 46 and ECR of ∼1160 μm. Across the cohort, B cells in HGG samples exhibited significantly greater CLF clustering intensity (mean CI = 92) over a shorter distance (mean ECR = 864 µm) than T cells (mean *CI* = 42, mean ECR = 1129 µm; Mann-Whitney U test, *P* = 2.69 × 10^-15^ and 3.57 × 10^-17^, respectively; **Fig. 5D-E**; **Supplementary Fig. S5F-K**). Moreover, radii estimated by use of the CLF are ∼4 to 8-fold greater than median radii for ICs obtained using KNN and DBSCAN clustering methods (∼280 & ∼140 µm, respectively). We therefore conclude that B and T cells in PCa are non-randomly distributed to a length of ∼1 mm, which lies well outside of the relatively compact structures that define the “cores” of BICs and TICs (**Fig. 2A-B, 3A, 4A-B**).

Antitumor immunity depends on cognate interactions between B and T cells. To quantify whole-slide co-clustering of T and B cells (**Fig. 5F-G**), we applied the multitype CLF (MCLF) which quantifies the spatial distribution of cell - cell interactions (**Methods, Supplementary Data S2, Supplementary Fig. S5D**). We observed a greater co-clustering radius in HGG compared with LGG (ECCR = 915 vs. 793 μm, Mann-Whitney U and Leven’s test, *P* = 0.049 & *P* = 0.001, **Fig. 5H, Supplementary Fig. S5L-M**). Thus, B and T cells are more likely to interact over longer spatial scales in HGG tumors, extending well beyond IC cores. These findings further confirm that HGG has a more spatially organized distribution of B and T than LGG and that non-random distributions of B and T cells can be detected at both short and long radii (from single cell diameters to *r* ∼1 mm), potentially reflecting a diversity of functional interactions. A strength of the MCLF in this setting is that it yields a summary statistic that encompasses features at the level of cell-cell interactions (spatial proximity) and mesoscale architectures (organization of an IC) across an entire specimen.

### Evidence of active immunosurveillance and potentially targetable T cell programs in PCa

Effective T cell-mediated immune surveillance requires antigen presentation by tumor cells and interaction with cytotoxic T cells (CTLs). Loss of HLA class I expression is a common route of escape from immune surveillance. We compared normalized HLA-A expression in AMACR-positive tumor cells with the matched AMACR-positive benign epithelium from the same sections (as a control) but found no significant difference (paired *t*-test, *P* = 0.39, **Supplementary Fig. S6A**). CTLs can be identified in CyCIF data based on expression of the cytolytic protease granzyme B (GZB) and we found that CD8^+^ PD-1^+^ GZB^+^ cells comprised up to 0.9% of all T cells in PCa specimens and were almost 2-fold more abundant in HGG than LGG PCa (median of total number of CTLs divided by all T cells of ∼3 × 10^-3^ and ∼4 × 10^-4^, respectively; Mann-Whitney U tests, *P* = 0.029, **Supplementary Fig. S6B**). The GZB^+^ fraction of CD8 T cells and CD8^+^ GZB^+^ of the T cells were significantly higher in HGG as compared to LGG (Mann-Whitney U test, *P* = 0.001 and *P* = 0.068, **Fig. 6A; Supplementary Fig. S6C**). When we scored the proximity of the latter to the tumor cells using a stringent interaction distance of ≤ 2 µm, we observed a significant difference between HGG compared to LGG PCa (Mann-Whitney U test, *P* = 0.006; **Fig. 6B-D**). These data suggest that HGG PCa contains CTLs, a subset of which are in proximity to tumor cells, and that these tumor cells express HLA-A and potentially other Class I molecules at normal levels. Thus, it seems likely that tumors, particularly in HGG localized PCa, are subject to active immune surveillance by T cells.

**Figure 6.**
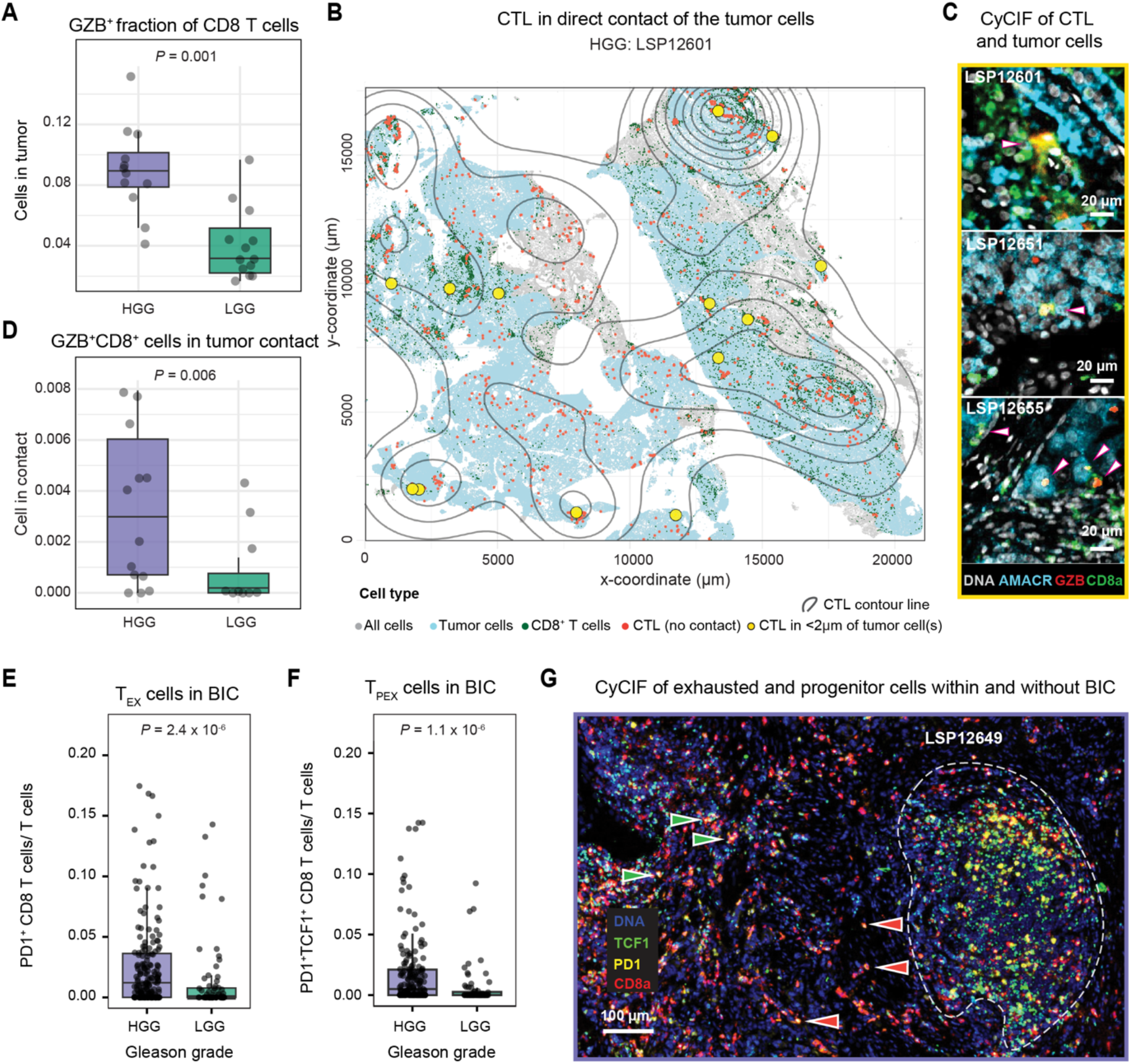
High-grade PCa exhibits spatially restricted enrichment of stem-like T cells and dendritic cells in B cell clusters. **A**, Cytotoxic T cell (CTL; GZB^+^ CD8^+^) proportions in the tumor for HGG and LGG samples (*P* value represents Mann-Whitney U). **B**, Pseudo-image of an HGG sample showing the tumor region, CD8^+^ T cells, and CTLs. CTLs in contact with tumor cells (<2 µm) are shown in yellow, with contour lines depicting the spatial frequency of CTLs. **C**, CyCIF images of CTLs in contact with tumor cells, indicated by overlaid arrows. **D**, Proportion of CTLs in contact with tumor cells (<2 µm) for HGG and LGG samples (*P* value represents Mann-Whitney U test). **E**, Percentage of T_EX_ exhausted CD8 T cells (PD-1^+^ CD8^+^) and **F**, T_pex_ progenitor-exhausted cells (TCF1^+^ PD-1^+^ CD8^+^) relative to total T cells (CD3d^+^) within BIC regions, stratified by high-grade (HGG, purple) and low-grade (LGG, green) tumors (*P* value represents Mann-Whitney U test). **G**, Representative CyCIF image highlighting increased exhaustion and progenitor-exhausted cell states adjacent to a BIC. TCF1^+^ PD-1^+^ CD8 T cells and TCF1^-^ PD-1^+^ CD8 T cells are indicated by right-pointing green arrows and left-pointing red arrows, respectively. The BIC area is encircled by a white dashed line.

TLSs have been shown, in some cancers, to play a role in responsiveness to PD-1 and PD-L1-directed immunotherapies(2,64). Whereas antigen-experienced CD8^+^ PD-1^+^ cells can be found both in TLSs and in the tumor compartment(58), PD-1^+^ TCF1^+^ CD8^+^ T cells (which likely corresponded to “precursor of exhausted” or T_pex_ cells)(58) are generally enriched in TLSs, where they interact with proliferating B cells (**Supplementary Fig. S6D)**. In our cohort, we found that CD8^+^ PD-1^+^ T cells comprised a ∼5-fold greater proportion of all T cells in BICs from HGG than LGG PCa (Mann-Whitney U test, *P* = 2.4 x10^-6^, **Fig. 6E; Supplementary Fig. S6E**) and that T_pex_ cells were ∼7-fold more prevalent in HGG BICs (Mann-Whitney U test, *P* = 1.1 × 10^− 6^, **Fig. 6F**). In contrast, no significant difference in the abundance of PD-1^+^ CD8^+^ cells was observed outside of BICs with or without stratification on TCF positivity (Mann-Whitney U test, TCF1^-^ : *P* = 0.11 and TCF1^+^: *P* = 0.19, **Fig. 6G**). These findings show that HGG prostate cancer exhibits significantly higher proportions of T_pex_ cells within BICs compared to LGG, highlighting the potential role of the TLS in supporting therapeutically targetable T cell programs.

## DISCUSSION

Both metastatic and primary PCa are often considered immunologically “cold”(65) cancers, in part because single-agent immune checkpoint blockade strategies have achieved only limited success in unstratified trials. However, we found that a substantial fraction (∼30-40%) of HGG PCa (Gleason grade groups 4 and 5) is as highly infiltrated with B cells as dMMR CRC cancer, an archetype of an immunologically hot cancer. T cells are ∼4-fold less abundant in PCa than dMMR CRC, but they are nonetheless organized into immune complexes with the hallmarks of an active immune response. A subset of effector CD8 T cells is also PD-1 positive, expresses GZB, and is in intimate contact with tumor cells, all features of anti-tumor cytotoxicity(66). The TCF1^+^ PD-1^+^ CD8^+^ cells present in HGG ICs are likely to represent precursor exhausted (T_pex_) T cells, which are increasingly recognized as critical for responsiveness to ICI therapies: T_pex_ cells retain proliferative potential and the ability to differentiate into effector cytotoxic T cells following PD-1/PD-L1 blockade(67,68). Thus, a subset of HGG PCa is not immunologically cold: it contains abundant B and T cell focused immune clusters, stem-like T cells, and CTLs in the tumor compartment itself.

A hallmark of an active immune response is coordination among diverse cell types through cell-cell signaling and spatial clustering. The best organized ICs act as hubs for antigen presentation, T cell co-stimulation, B cell activation and affinity maturation, as well as other local anti-tumor responses(69). In multiple tumor types(1) the presence of TLSs(2,57,64,70,71) is associated with improved clinical outcomes as well as benefit from immunotherapy(2,64). Prior H&E and IHC studies have suggested that immune cells in PCa are organized spatially, but the absence of an objective means of identifying and classifying ICs has impeded analysis. We find that classical germinal center morphology, as judged by trained histopathologists from H&E images(72,73), is present on a minority of ICs in PCa, but that many ICs lacking classical morphology have immunological features associated with TLSs including clusters of follicular dendritic cells, proliferating B cells, and stem-like T cells. We therefore combined statistical methods to quantify immune cell distributions and their relationships to tumor grade; these approaches are likely to be broadly useful for other cancer types and biological variables and we therefore make them available in a single R package (*tlsR*).

Although prior H&E and IHC studies have suggested that immune cells in PCa are spatially organized, the absence of an objective way to parameterize immune clusters (ICs) has impeded analysis. To address this, we developed a set of complementary statistical tools in the tlsR R package: KNN and DBSCAN clustering to identify immune clusters (**Fig. 2B, Fig. 4D, Supplementary Fig. S7A**), a multitype Ripley’s L-function to test for non-random organization apart from discrete clustering (**Fig. 5A**), and ICAT to quantify cluster geometric organization (**Fig. 3A-B**). The strength of ICAT in this context, relative to more complex deep learning methods(74), is that ICAT is a geometrical property of an IC or a specimen that can be interpreted in terms of familiar concepts such as shape and compactness. Together these approaches show that ICs are both more prevalent and more organized in high-grade than low-grade PCa, that B and T cells co-cluster at radii up to ∼1 mm, and that the best-organized ICs carry the immunological hallmarks of mature TLSs, including follicular dendritic cells, proliferating B cells, and stem-like T cells. However, only a minority of these ICs had classical germinal center morphology in H&E images suggesting that this is too strict a criterion for ICs in which B and T cells have the potential to interact functionally(44).

Despite the presence of immune cells in the tumor microenvironment (TME) of PCa, immune checkpoint inhibition (ICI) trials have been underwhelming in mCRPC(75,76) and there is no established role for tumor immune profiling in guiding treatment. Furthermore, the role of immunotherapy for the treatment of primary high-risk PCa remains poorly understood. Tumor-infiltrating CD4^+^ and CD8^+^ T cells in primary PCa have previously been shown to express the immune checkpoint marker and ICI target PD-1(77). However, bulk RNA-seq studies have suggested that prostate tumors carry lower T cell fractions and higher macrophage and neutrophil signatures than adjacent benign tissue(78). Moreover, single-cell RNA-seq (scRNA-seq) surveys of primary PCa have suggested the presence of an immune-suppressive microenvironment rich in macrophages and inflammatory monocytes with most CD8^+^ T in an exhausted state(79,80). Despite this, multiple novel immunotherapy approaches being evaluated for the treatment of mCRPC. That some of these approaches are demonstrating clinical activity suggests that immunotherapy can be effective in the right patient population, particularly earlier in the disease spectrum, such as high-risk treatment-naïve localized PCa. One possibility is that selection for tumors enriched in CD8^+^ T cell infiltration, as currently being explored for mHSPC(81), or for tumors with organized ICs and T_pex_ cells (as in this study) will be sufficient to uncover a subset of ICI-responsive patients missed in unstratified trials. Therapeutic approaches that focus on mobilizing B cells or activating T cells by mechanisms that do not involve inhibition of PD-1-PD-L1 binding are an alternative possibility.

This study has several limitations. First, the cohort size was modest (n=29), limiting statistical power. Future studies using higher-throughput spatial imaging platforms(45) would enable evaluation of substantially larger cohorts. Second, our analyses were performed exclusively in radical prostatectomy specimens. In contrast, clinical risk stratification and Gleason grading in localized disease relies primarily on diagnostic needle biopsies that may not reliably capture TLSs or broader immune architecture. Nonetheless, our observation that TLS presence strongly correlates with overall immune infiltration suggests that immune profiling of biopsies may still identify highly inflamed tumors. Third, our study focused only on hormone-sensitive PCa. In mCRPC, where therapeutic options are more limited, tumors often exhibit a more immunosuppressive microenvironment and low response rates to immune checkpoint blockade, even in the setting of immune infiltration(82). Whether a subset of mCRPC as defined by enriched ICs that may be more responsive to immunotherapy remains unknown and warrants further investigation.

## ACKNOWLEDGEMENTS

This work was supported primarily by an ASPIRE Award from The Mark Foundation for Cancer Research and by Ludwig Cancer Research; additional support was provided by NCI grants P50CA180995 and R01 CA273914-01 (AP), Michael & Lori Milken Family Foundation-Prostate Cancer Foundation Challenge Award (AP), V Foundation Translational Cancer Research Award (AP), U01-CA284207 and R50-CA252138 to ZM, Finnish Cultural Foundation Grant No. 211288 (Ali. A), Wong Family Foundation Award (JW), ASCO Young Investigator Award (JW), Prostate Cancer Foundation Young Investigator Award (JW) and NCI K99CA300495 (JW); S.C. was supported by training grant T32-CA009216 (NCI) and the HMS Program in Neuroscience Award. Histopathology support was provided by P30CA06516; additional support was provided by P50-CA272390, P01-CA228696, DOD W81XWH-21-PCRP-DSA, and DOD HT94252410415.

## AUTHOR CONTRIBUTION STATEMENTS

A.A., J.A.W., J.L., A.P., and P.K.S. conceived and designed the study. A.P. and P.K.S. supervised and acquired the funding. A.A. and J.A.W. developed and implemented the data analysis workflow, produced the figures, and interpreted the results. J.L. and B.K. acquired the images and performed initial image analysis. A.G., F.S., and S.C. provided histopathological annotations. A.A., J.A.W., and P.K.S. wrote the manuscript with input from all authors.

---

**Supplementary Figure 1.**
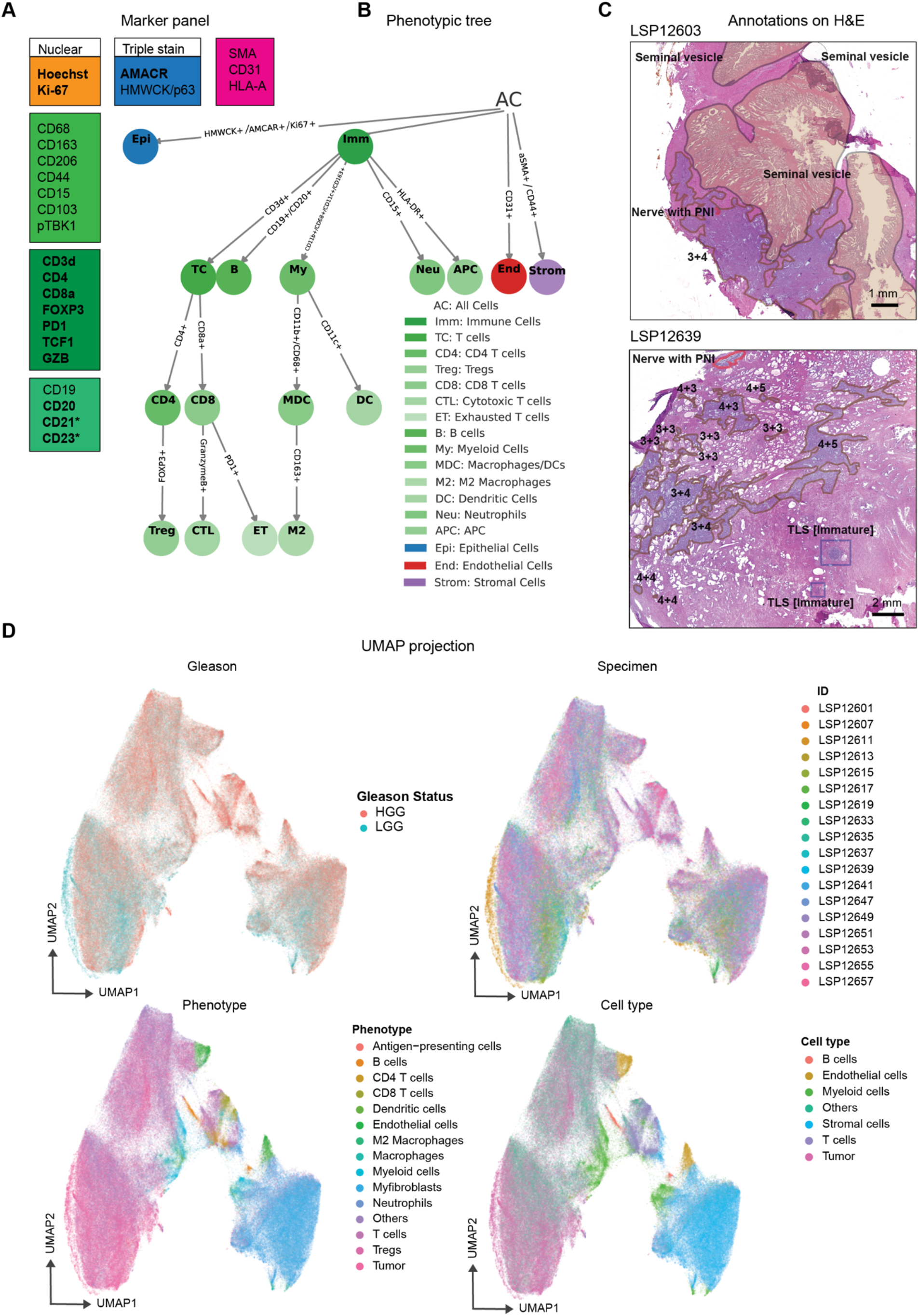
Cell phenotyping and quality control. **A**, CyCIF markers grouped by function: epithelial markers (blue), stromal markers (pink), T-cell markers (red), innate-immune markers (purple), B-cell markers (green), and nuclear markers (orange). Key markers for this study are shown in bold and markers used specifically to study follicular dendritic cells in a subset of specimens are marked with an asterisk. **B**, Phenotype hierarchy derived from the full marker set as used to assign cell types and states. **C**, Examples of genitourinary (GU) pathologist annotations on H&E sections; two pathologists independently annotated the specimens, and their annotations were found to be in excellent agreement. **D**, UMAP embedding of balanced sampling of 10,000 tumor and stromal cells per patient based on scaled, ungated marker intensities; points are colored by Gleason status, cell phenotype, and patient ID.

**Supplementary Figure 2.**
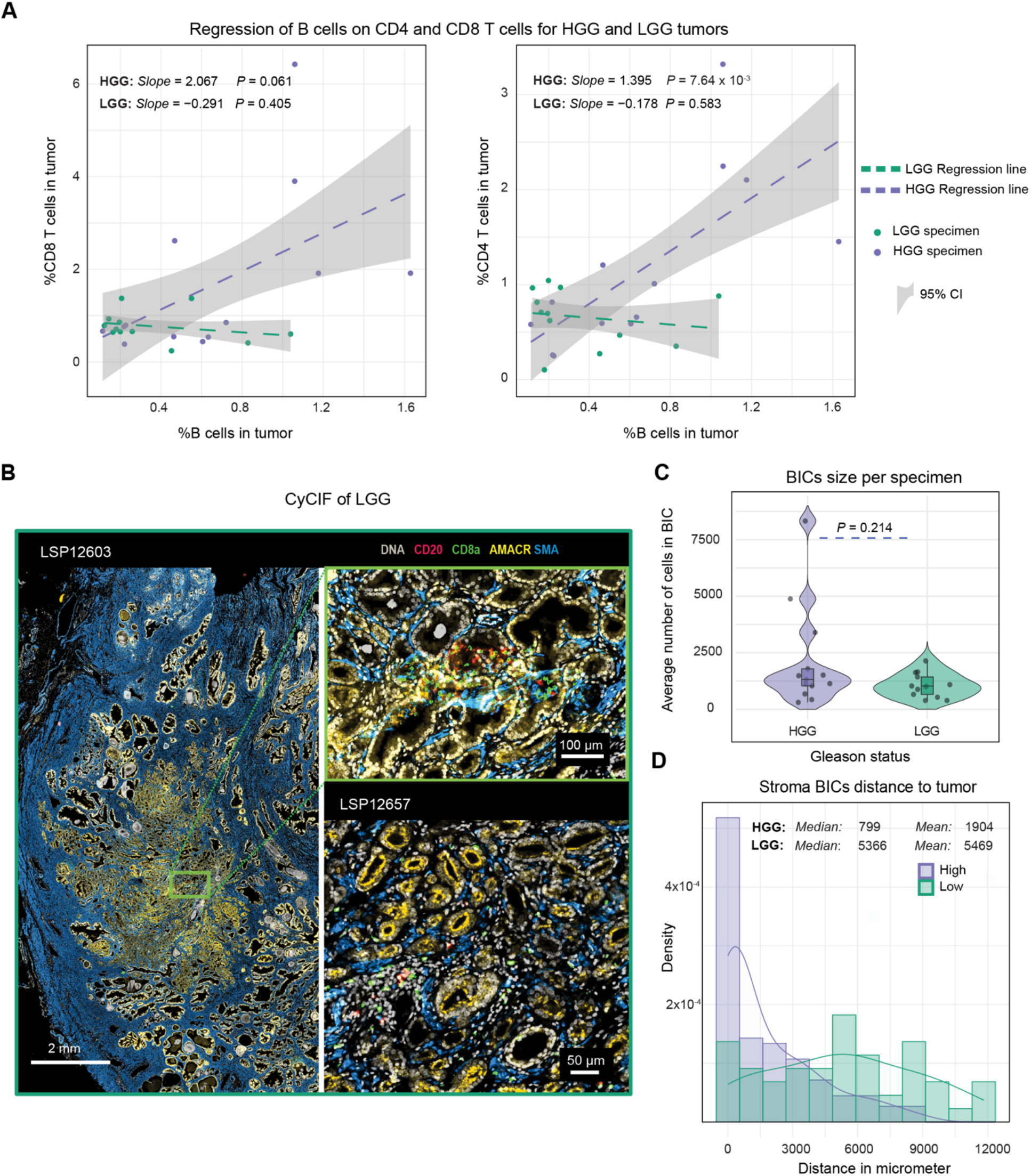
B and T cells densities and BICs exhibit distinct patterns between low -grade and high-grade PCa. **A**, Regression fit and 95% confidence intervals of the proportion of intratumoral CD8^+^ (left) and CD4^+^ (right) T cells compared with CD20^+^ cells by Gleason grade. Annotated values represent the *P* value of the fit (*F*-test) and the maximum likelihood estimate of the slopes. **B**, Whole slide CyCIF image of a LGG case displayed with markers for nuclear DNA, smooth-muscle actin (SMA), CD8. Two zoomed-in exemplar regions are shown to the right. **C**, Mean number of cells per BIC per tumor stratified by tumor grade, with *P* value representing Mann Whitney U test. **D**, Histogram and smoothed curve of the tumor-BIC distances for extra-tumoral BICs, by tumor grade group.

**Supplementary Figure 3.**
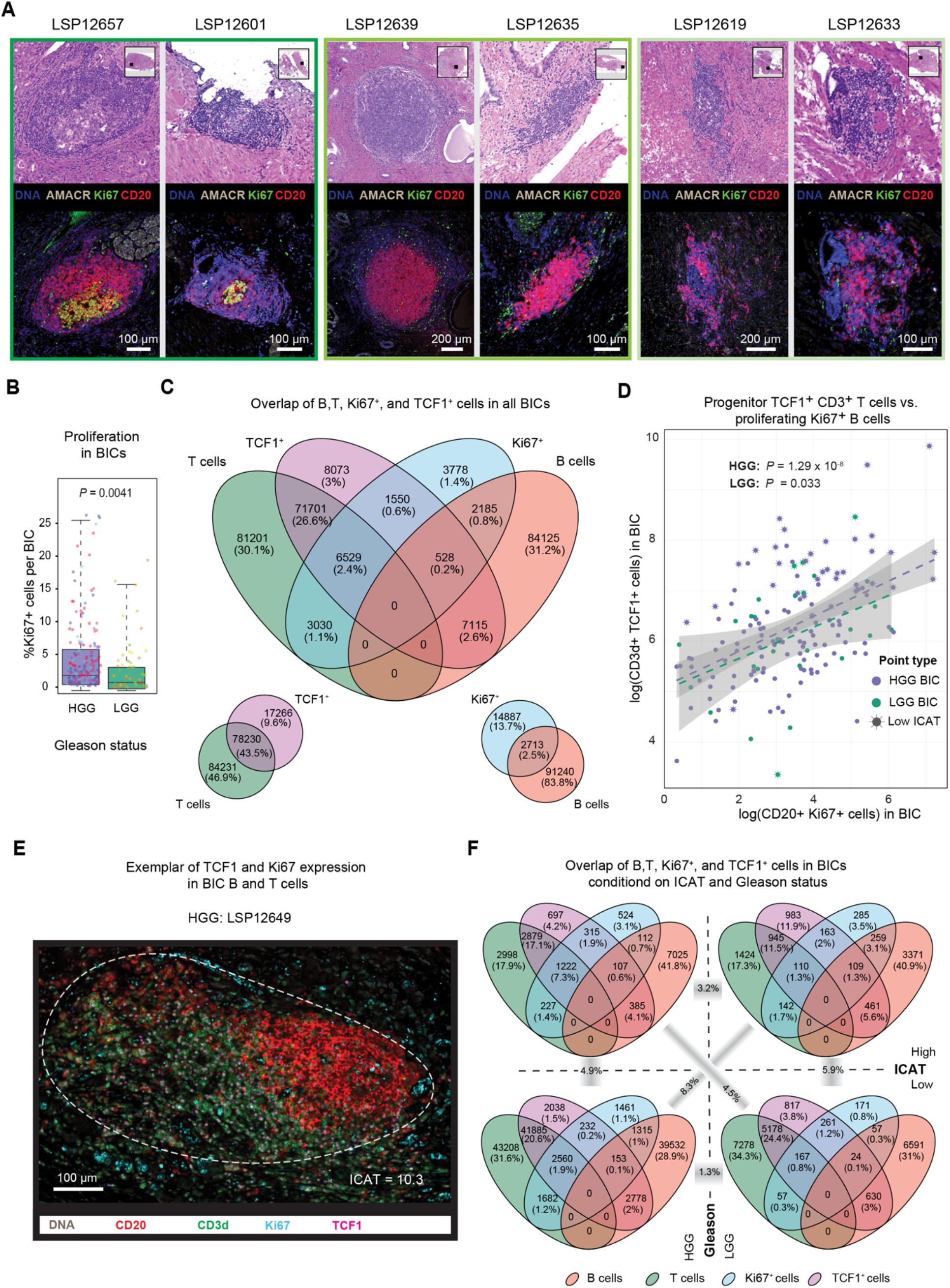
Spatially organized BICs contain Ki67^+^ proliferative cells and TCF1^+^ T cells. **A**, Examples of BICs with mature TLS morphology (left), immature TLS morphology (middle) and disorganized morphology (right) based on expert H&E annotation and same-section identical magnification CyCIF. **B**, Proportion of BIC cells that are Ki67^+^, colored by specimens and stratified by tumor grade (*P* value represents Mann-Whitney U test). **C**, Venn diagram across all BICs showing the respective number and proportion of the cells for CD20^+^ (B cells), CD3d^+^ (T cells), TCF1^+^, and Ki67^+^. **D**, The regression line and 95% confidence interval between the logarithmic scale of Ki67^+^ B and TCF1^+^ T cells within BICs, stratified by tumor grade. BICs with an ICAT in the lowest quartile (most organized; ICAT < 13.7) are shown with a star. *P* value represents *F*-test of the regression fit with *R*^2^_HGG_ = 0.22 and *R*^2^_LGG_ = 0.16. **E**, CyCIF image depicting Ki67 and TCF1 signal on B and T cells in a BIC (encircled in white dashed line). **F**, Venn diagrams in (**C**) stratified by ICAT score (top to bottom) and Gleason grade (left to right). The cross dissimilarities between each diagram’s proportion vector are measured by Jensen Shannon Divergence (JSD) reflected for the relevant pairs.

**Supplementary Figure 4.**
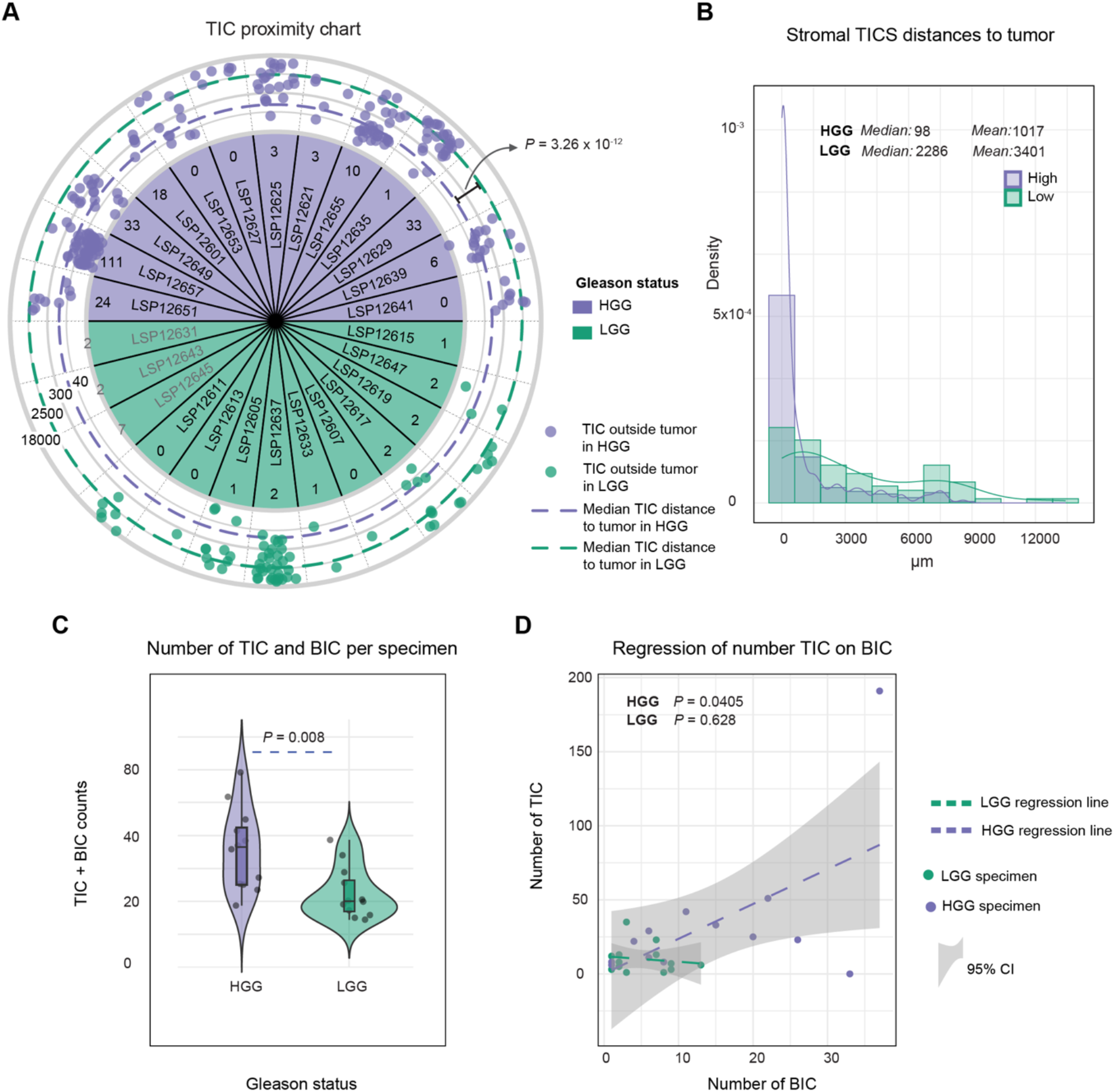
T cell immune clusters are more numerous and more spatially proximal to tumors in high-grade PCa. **A**, Schematic depiction of the logarithmic distance from each T cell immune cluster (TIC) to the tumor compartment for extra-tumoral TICs (outer rings in μm), and the total count of intra-tumoral TICs for intratumoral TICs (inner ring). *P* value represents the Mann-Whitney U test of the TIC-tumor distance comparing HGG and LGG PCa. Three samples without annotated tumor regions on our slide are highlighted in grey. **B**, Histogram and smoothed curve of the tumor-TIC distances for extra-tumoral BICs, by tumor grade group. **C**, Total number of BICs and TICs in each specimen stratified by tumor grade (*P* value represents Mann-Whitney U test). **D**, Regression fit and 95% confidence interval for the number of TICs compared with number of BICs, stratified by tumor grade (*P* values represent *F*-test of the fit).

**Supplementary Figure 5.**
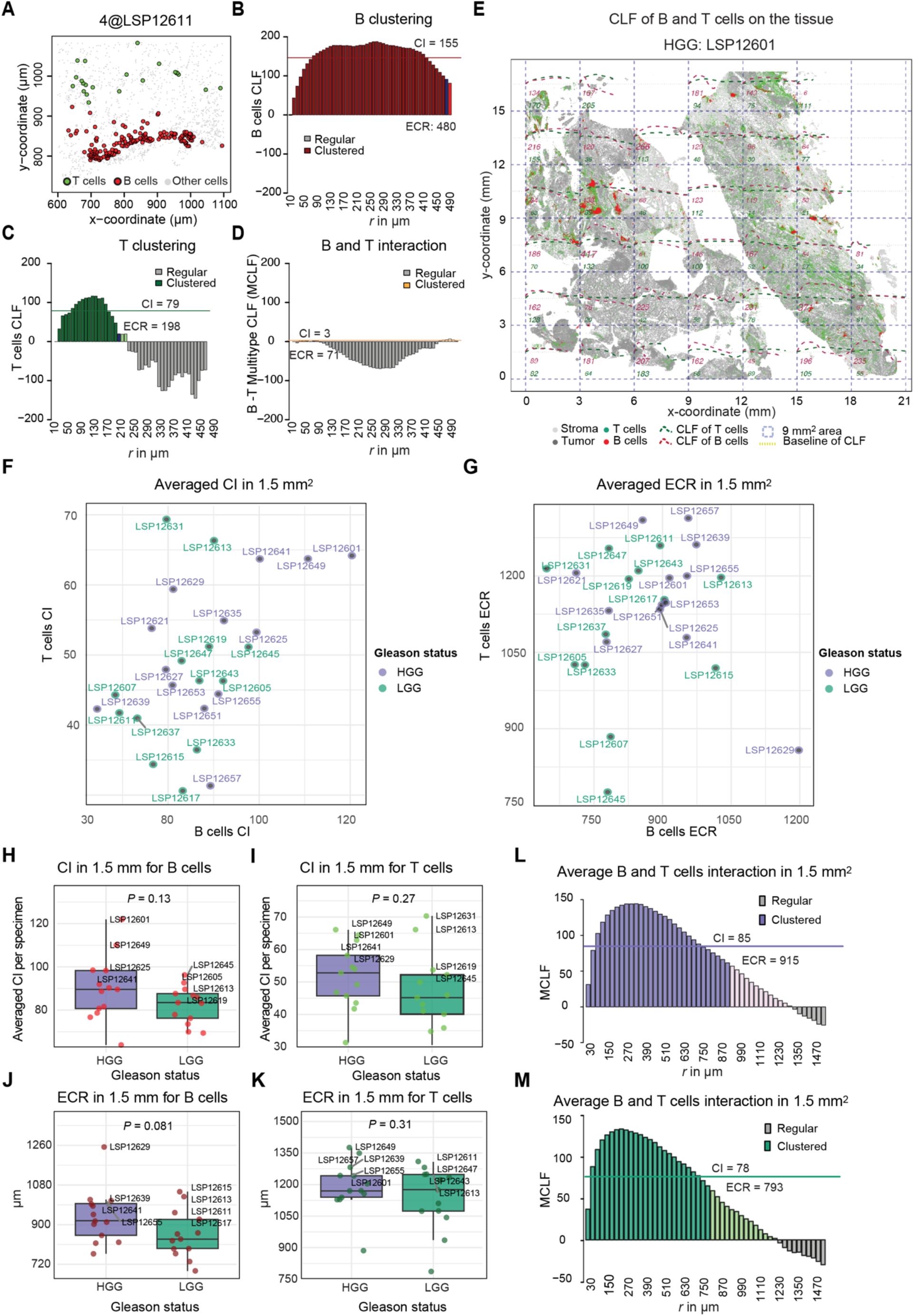
The spatial interaction of B and T cells across whole-slide prostate cancer images. **A**, Pseudo-image (cells are circles) showing B and T cell communities on the tissue in a LGG case. **B**, The corresponding centered L-function (CLF) for B cells, highlighting a consistently higher clustering intensity (CI) over a larger radius (ECR). **C**, The CLF for T cells, indicating a phase shift toward complete spatial randomness (CSR) and regularity, with lower CI and ECR. **D**, The interaction between B and T cells, characterized by the multitype CLF (MCLF), underscoring consistent regularity indicative of complete segregation. **E**, Depiction of whole-tissue B (CD20^+^) and T (CD3d^+^) cells, along with a clustering survey using the centered L-function (CLF) over 9 mm^2^ square areas, binned over increasing 30-μm radii (dashed curves), with marked positive integral values in red and green for each area, respectively. **F**, Point plot of clustering intensity (CI) for B and T cells, stratified by Gleason status, with annotations for specimen identifiers. **G**, Point plot of effective clustering radius (ECR) for B and T cells, stratified by Gleason status, with annotations for specimen identifiers. **H**, CI for B cells, stratified by Gleason status, showing respective comparison Mann-Whitney U test *P* values, with annotations for the four highest values in each plot including specimen identifiers. **I**, Similar to (**H**), boxplots for T cells. **J**, Similar to (**H**) boxplots for ECR. **K**, Similar to (**H**) boxplots for T cells and ECR. **L**, Multitype CLF (MCLF) depicting the interaction between B and T cells, integrated across all fourfold systematic bootstrapped 9 mm^2^ square areas for each specimen and averaged for each 30-μm radius bin in HGG samples, overlaid with co-clustering intensity (CCI) and estimated co-clustering radius (ECCR). **M**, Multitype CLF (MCLF) depicting the interaction between B and T cells, integrated across all fourfold systematic bootstrapped 9 mm^2^ square areas for each specimen and averaged for each 30-μm radius bin in LGG samples, overlaid with co-clustering intensity (CCI) and estimated co-clustering radius (ECCR).

**Supplementary Figure 6.**
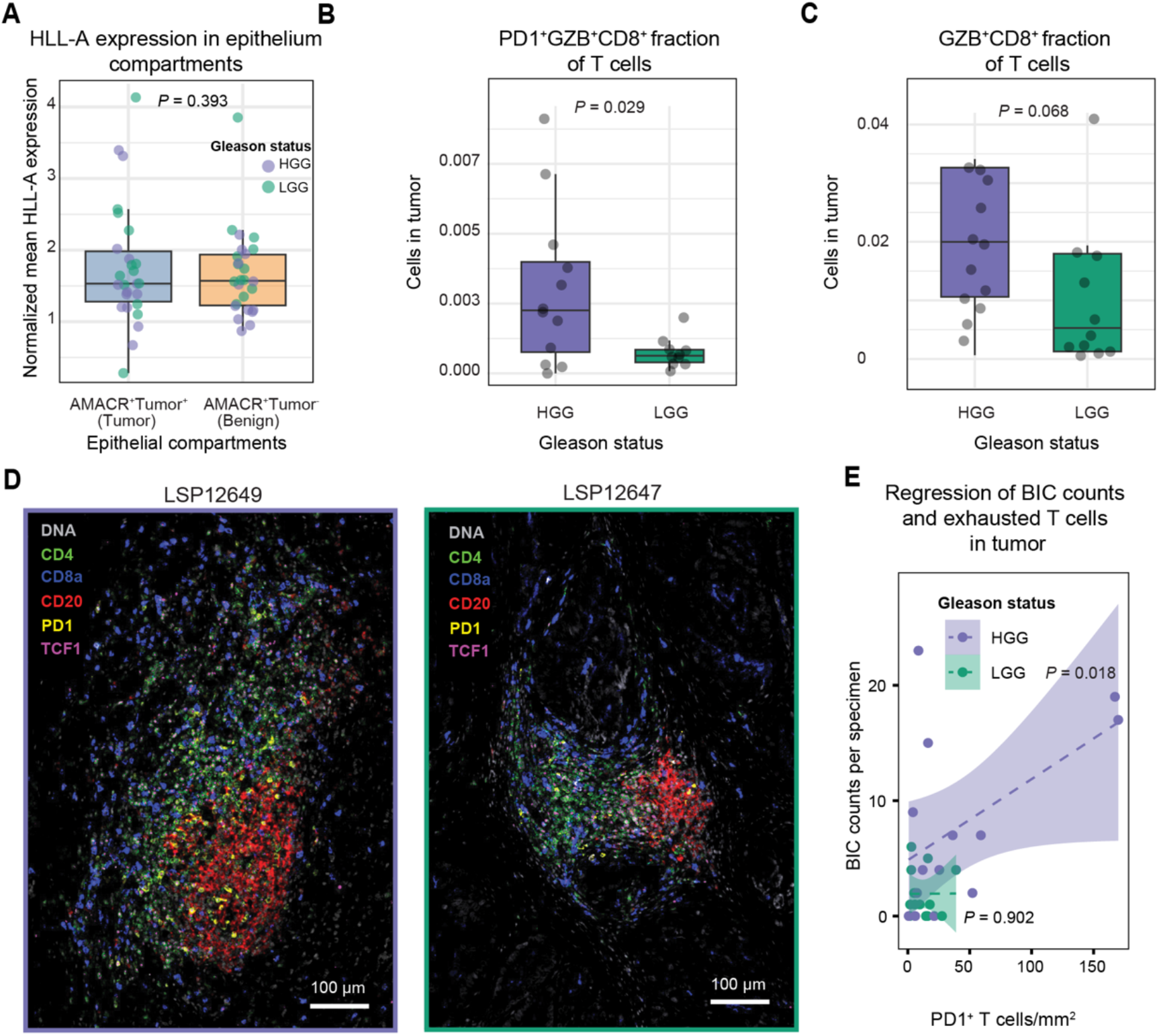
Further quantification of T cell subtypes and HLA-A expression in HGG versus LGG PCa. **A**, Mean normalized HLA-H immunofluorescence expression in cells from the non-tumor and tumor compartments of the tissue (*P* value represents paired *t*-test). **B**, Proportion of PD-1^+^ GZB^+^ CD8^+^ cells among T cells in the tumor for HGG and LGG tumors (*P* value represents Mann-Whitney U test). **C**, Proportion of GZB^+^ CD8^+^ cells among T cells in the tumor for HGG and LGG tumors (*P* value represents Mann-Whitney U test). **D**, Portion of CyCIF images denoting increased expression of PD-1 and TCF1 around the BIC in two HGG specimens. **E**, Regression fits of the number of BICs in each specimen against the density of PD-1^+^ T cells in the tumor, stratified by HGG and LGG tumors, with corresponding *F*-test *P* values for the fits and 95% confidence intervals.

**Supplementary Figure 7.**
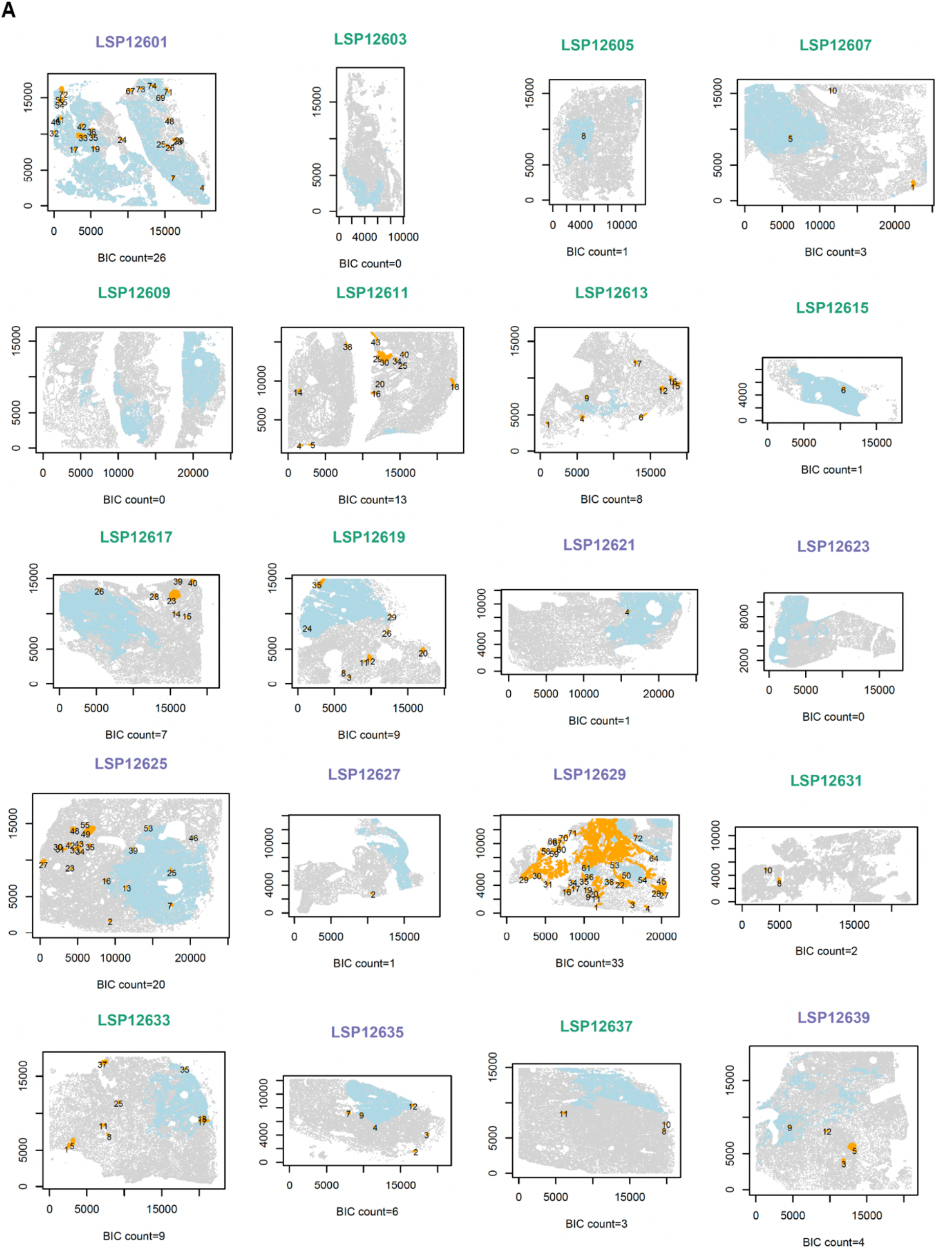
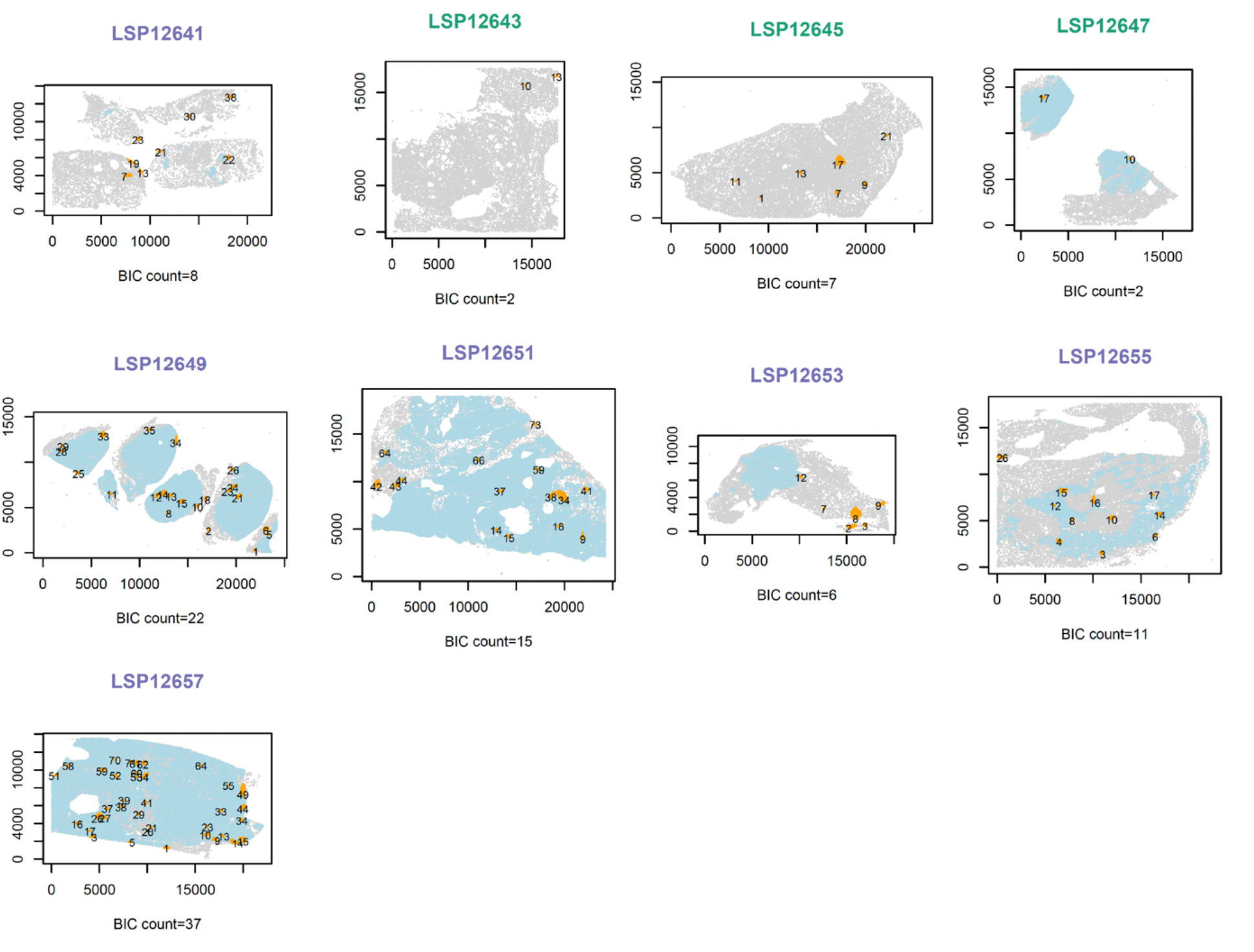
Tumor and BIC annotations across all samples. **A**, Pseudo-images (cells are circles) of the whole-slide tissue annotated with the tumor areas (light blue) and numbered detected BICs in orange with their identifier. Titles stratified for HGG and LGG specimen with purple and green, respectively and the axes denote the coordinates in μm.

## Supplementary Data File

### 1. Independent component analysis of the immune clusters

The goal is to find the aggregated standard deviation of the points in the x-y space per cells in a given immune cluster (IC) by prioritizing the canonical shapes. Note that we can use the Independent Component Analysis (ICA) matrix decomposition for cells in IC **X** as

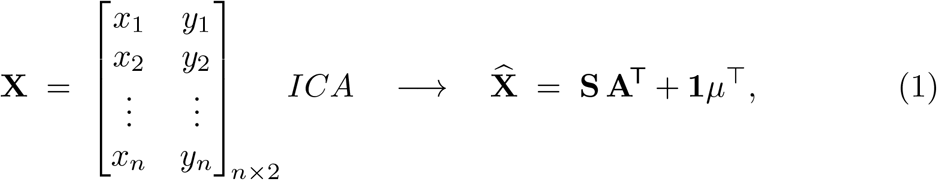

where **S** = [**s**_1_ **s**_2_]_*n*×2_ are the, **A** ∈ ℝ^2×2^ is the **mixing matrix, 1** ∈ ℝ^*n*^ is the all-ones vector, and 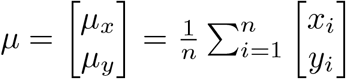.

The 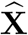 space is the ‘best’ independent representation of the X in the same unit space

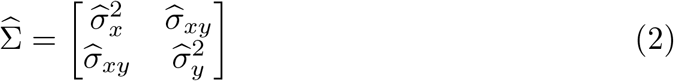

as the covariance matrix with trace and determinant of

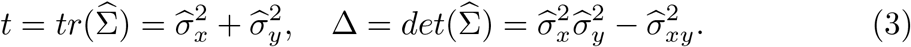

Now the standard deviation matrix is given by

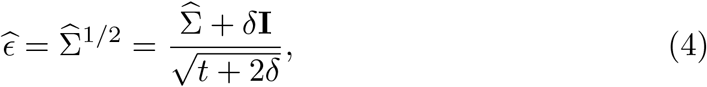

where 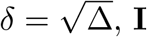 is the identity matrix, and 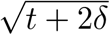 is

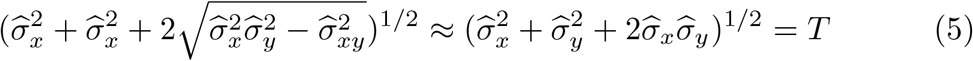

which is so as under statistical independence assumption of 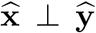, we have 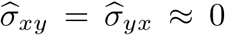 tacitly penalizing cases where correlation between **x**^′^ and **y**^′^ still persist underlying point configurations with aberrant non-collinear shapes that independent components are not perfectly achievable. This leads to 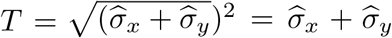 which is exactly the trace of the standard deviation matrix 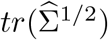.

So the per-cell projected (percentage if *α* = 100) and normalized total standard deviation of a cluster based on the underlying independent coordinates is

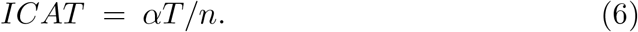

### 2. Ripley *K*-, *L*-functions and derived clustering indices

Let’s define the set of B and T cells with the following notations:

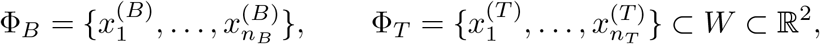

with |*W*| the tissue area and *λ*_*B*_ = *n*_*B*_*/* |*W*|, *λ*_*T*_ = *n*_*T*_ */* |*W*| the global intensities of these cells.

#### Theoretical *K*-functions

For a single cell type *i* ∈ {*B, T*} the theoretical *K*-function as the averaged intensity of given cells for a set radius *r* is presented by:

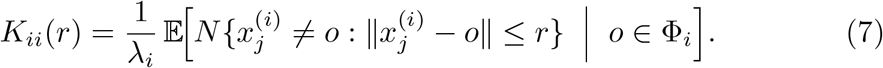

Similarly, the averaged cross-type interaction of cells of type *j* ∈ *T*, within a given radius *r* of type *i* ∈ *B* is given by:

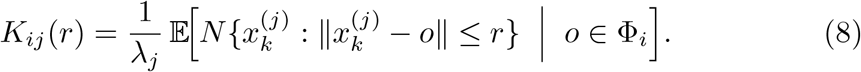

#### Edge-corrected estimator

Now in attempt to estimate theoretical functions the edge-corrected generic estimator is:

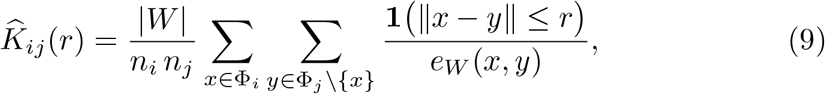

where **1**(∥*x* − *y*∥ ≤ *r*) = 1 when the inter-cell distance does not exceed *r*, and *e*_*W*_ (*x, y*) is Ripley’s isotropic edge weight.

#### Variance-stabilized and centralized *L*-function

The above function allows us to derive two more intuitive functions formally referred to as variance stabilized (by taking the square root and scaling for the constant *π*) and centered (by further removing the increasing effect of *r*) *L*-functions as below:

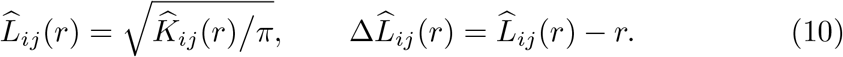

For much of the analysis, we rely on the latter formula (referred to as CLF in the manuscript) as the estimate of the cells dispersion patterns with 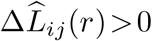 signaling attraction, = 0 CSR, and *<* 0 as inhibition.

#### Summarizing the CLF

Based on the above conventional formulae, here we define three novel indices to capture relevant dimensions of this function as clustering intensity, half-mass radius, and effective clustering radius. Before that lets define the positive envelope 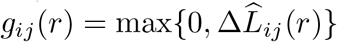 and denote by 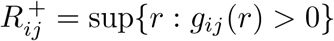 the largest radius at which any positive deviation persists.

##### 1. Clustering intensity

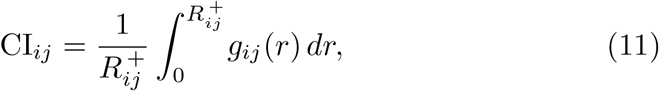

i.e. the mean magnitude of attraction across the range on which it is present. Since the *R* ^+^ is specific to each window, this quantity simply is scaling the integral of the positive (attraction) portion of the CLF.

##### 2. Half-mass radius

We also define an instrumental *R*50 index as radius enclosing first-half of the total positive clustering mass by

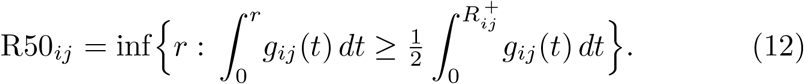

##### 3. Effective clustering radius

Since in majority of the cases the CLF is right-skewed, harboring on the previous index allows us to define

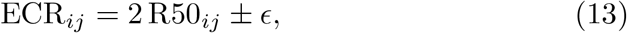

which is providing an operational outer limit of attraction that remains slightly inside the point where 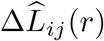 first returns to the CSR baseline, with *ϵ* as half of the binning used for *r* toward estimation of the *K*-function. This is a conservative estimation of the clustering radius statistically below the radius on which the CLF passes through the CSR confidence interval.

Together, CI_*ij*_, R50_*ij*_, and ECR_*ij*_ compress the full centred *L*-curves into three interpretable numbers describing, respectively, the average strength, the core scale, and the effective spatial extent of the cells interactions. More precisely the B-, T-, and B-T cells interactions are obtained when *i* = *j* ∈ *B, i* = *j* ∈ *T*, and (*i, j*) ∈ (*B, T*), respectively.

**Type** Package

## Package ‘tlsR’

May 8, 2026

**Type** Package

**Title** Detection and Spatial Analysis of Tertiary Lymphoid Structures

**Version** 0.3.0

**Date** 2026-04-02

**Description** Fast, reproducible detection and quantitative analysis of tertiary lymphoid structures (TLS) in multiplexed tissue imaging.

Implements Independent Component Analysis Trace (ICAT) index, local Ripley’s K scanning, automated K Nearest Neighbor (KNN)-based TLS detection, and T-cell clusters identification as described in

Amiryousefi et al. (2025) <doi:10.1101/2025.09.21.677465>.

**License** MIT + file LICENSE

**URL** https://github.com/labsyspharm/tlsR

**Depends** R (>= 4.0.0)

**Imports** dbscan (>= 1.1-10), fastICA (>= 1.2-3), FNN (>= 1.1.3),

spatstat.explore (>= 3.0-0), spatstat.geom (>= 3.0-0), ggplot2 (>= 3.4.0), rlang (>= 1.0.0), grDevices, graphics, stats, methods

**Suggests** knitr, rmarkdown, testthat (>= 3.0.0)

**Encoding** UTF-8 **LazyData** true **RoxygenNote** 7.3.3 **VignetteBuilder** knitr **Config/testthat/edition** 3 **NeedsCompilation** no

**Author** Ali Amiryousefi [aut, cre] (ORCID:

Jeremiah Wala [aut] (ORCID:), Peter Sorger [ctb] (ORCID:)

**Maintainer** Ali Amiryousefi

**Repository** CRAN

**Date/Publication** 2026-04-23 20:40:02 UTC

2 *tlsR-package*

### Description

Fast, reproducible detection and quantitative analysis of tertiary lymphoid structures (TLS) in multiplexed tissue imaging data.

### Typical workflow

1. Load or prepare a named list of data frames (`ldata`), one per tissue sample. Each data frame must contain columns `x, y` (spatial coordinates in microns), and `phenotype` (character: “**B cell” / “T cell**” / other).
2. Run `detect_TLS` to label B+T co-localised regions.
3. (Optional) Run `scan_clustering` to identify windows of significant immune clustering via local Ripley’s L.
4. Run `calc_icat` to score the internal linearity/organisation of each detected TLS.
5. Run `detect_tic` to identify T-cell clusters outside TLS.
6. Use `summarize_TLS` to obtain a tidy summary table.
7. Use `plot_TLS` to produce publication-ready spatial plots.

### Author(s)

**Maintainer**: Ali Amiryousefi (ORCID) Authors:

- Jeremiah Wala (ORCID)

Other contributors:

- Peter Sorger (ORCID) [contributor]

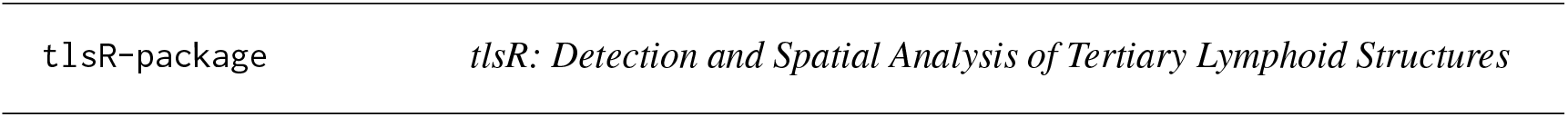

|  |  |
| --- | --- |
| tlsR-package | *tlsR: Detection and Spatial Analysis of Tertiary Lymphoid Structures* |

### References

Amiryousefi et al. (2025) doi:10.1101/2025.09.21.677465

*calc_icat* 3

**See Also**

Useful links:

- https://github.com/labsyspharm/tlsR

|  |  |
| --- | --- |
| calc_icat | *Calculate ICAT (Independent Component Analysis Trace) Index for TLS* |

### Description

Quantifies the spatial spread and linear organisation of cells within a detected TLS. FastICA is applied to the (x, y) coordinates of TLS cells to estimate independent components; the mixing matrix is used to reconstruct the data, and the ICAT index is defined as the normalised trace-standard-deviation of the reconstructed coordinates.

The index is always non-negative because it measures the average spatial spread per cell rather than the signed trace of the mixing matrix (which can be negative due to ICA sign ambiguity). Higher values indicate a more spatially extended, structured cluster.

### Usage

`calc_icat(patientID, tlsID, ldata = NULL`)

### Arguments

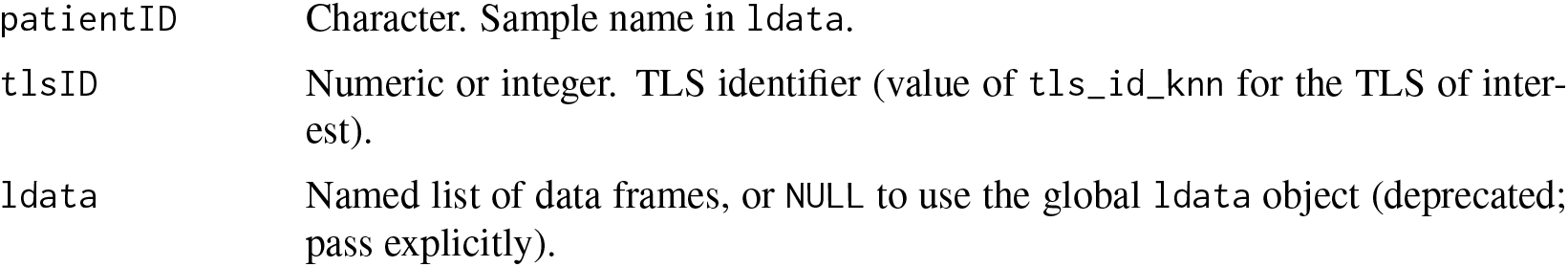

### Details

The ICAT index is computed as follows:

1. Centre the (x, y) coordinates of TLS cells.
2. Run `fastICA` with 2 components.
3. Reconstruct 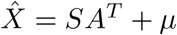.
4. Let *v*_1_, *v*_2_ be the marginal variances of 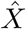.
5. 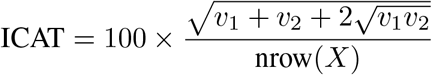

If the requested TLS contains fewer than 3 cells, or FastICA does not converge, the function returns `NA_real_` with an informative message rather than throwing an error.

### Value

A single non-negative numeric value (the ICAT index), or `NA_real_` if computation is not possible (fewer than 3 cells, or FastICA did not converge).

### Examples

~~~
data(toy_ldata)
ldata <-detect_TLS(“ToySample”, k = 30, ldata = toy_ldata)
if (max(ldata[[“ToySample”]]$tls_id_knn, na.rm = TRUE) > 0) {
    icat <-calc_icat(“ToySample”, tlsID = 1, ldata = ldata)
    print(icat)
}
~~~

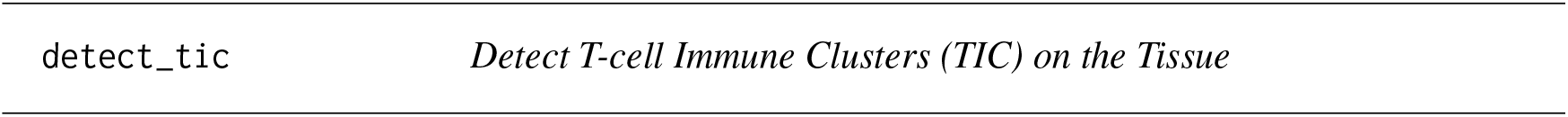

### Description

Applies HDBSCAN to T cells that lie outside of previously detected TLS regions to identify spatially compact T-cell clusters (TIC). Phenotype labels `“T cell”` and `“T cells”` are both accepted.

### Usage

`detect_tic(sample, min_pts = 10L, min_cluster_size = 10L, ldata = NULL)`

### Arguments

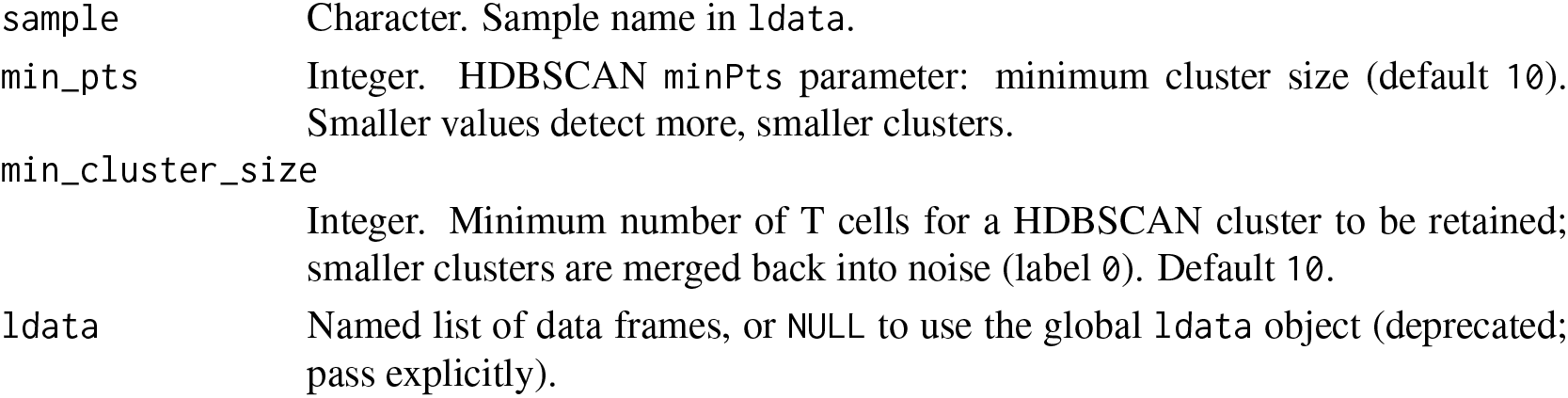

|  |  |
| --- | --- |
| sample | Character. Sample name in ldata. |
| min_pts | Integer. HDBSCAN minPts parameter: minimum cluster size (default 10). Smaller values detect more, smaller clusters. |
| min_cluster_size | Integer. Minimum number of T cells for a HDBSCAN cluster to be retained; smaller clusters are merged back into noise (label 0). Default 10. |
| ldata | Named list of data frames, or NULL to use the global ldata object (deprecated; pass explicitly). |

### Value

The input `ldata` list with the sample data frame augmented by one new column:

`tcell_cluster_hdbscan` Integer. 0 = noise / not a T-cell cluster; positive integer = TIC cluster ID. Non-T-cell rows receive NA.

### Examples

~~~
data(toy_ldata)
ldata <-detect_TLS(“ToySample”, k = 30, ldata = toy_ldata)
ldata <-detect_tic(“ToySample”, ldata = ldata)
table(ldata[[“ToySample”]]$tcell_cluster_hdbscan, useNA = “ifany”)
~~~

|  |  |
| --- | --- |
| detect_TLS | *Detect Tertiary Lymphoid Structures using a KNN-density approach* |

### Description

Identifies TLS candidates by finding regions of high local B-cell density that also contain a sufficient number of nearby T cells (B+T co-localisation). Phenotype labels `“B cell”` and `“B cells”` (and their T-cell equivalents) are both accepted.

### Usage

~~~
detect_TLS(
   LSP,
   ldata,
   k = 30L,
   bcell_density_threshold = 10,
   min_B_cells = 50L,
   min_T_cells_nearby = 10L,
   max_distance_T = 50,
   expand_distance = 80
)
~~~

### Arguments

|  |  |
| --- | --- |
| LSP | Character. Sample name in ldata. |
| k | Integer. Number of nearest neighbours used for density estimation (default 30, calibrated for 0.325 um/px imaging). |
| bcell_density_threshold | Numeric. Minimum average 1/k-distance (in microns) for a B cell to be considered locally dense (default 15). |
| min_B_cells | Integer. Minimum B cells per candidate TLS cluster (default 50). |
| min_T_cells_nearby | Integer. Minimum T cells within max_distance_T microns of the candidate cluster centre (default 30). |
| max_distance_T | Numeric. Search radius (microns) for T-cell proximity check (default 50). |
| expand_distance | Integer. The extended values from the boundary of the detected B-cells clusters that the T cells are being integrated (default 80). |
| ldata | Named list of data frames, or NULL to use the global ldata object (deprecated; pass explicitly). |

### Value

The similarly formatted `ldata` list, with the data frame for LSP augmented by three new columns:

`tls_id_knn` Integer. 0 = non-TLS cell; positive integer = TLS cluster ID.

`tls_center_x` Numeric. X coordinate of the TLS centre for TLS cells; `NA` otherwise.

`tls_center_y` Numeric. Y coordinate of the TLS centre for TLS cells; `NA` otherwise.

### Examples

~~~
# Use a 70% sample of the data to keep CRAN check time under 10s. # TLS detection requires sufficient cell density; 70% preserves
# the spatial structure needed for reliable detection.
# For production use, run on the full dataset (see \donttest{} below). data(toy_ldata)
set.seed(42)
idx <-sample(nrow(toy_ldata[[“ToySample”]]),
         size = floor(0.7 * nrow(toy_ldata[[“ToySample”]])))
sub_ldata <-list(ToySample = toy_ldata[[“ToySample”]][idx,])
ldata <-detect_TLS(“ToySample”, k = 30, ldata = sub_ldata)
table(ldata[[“ToySample”]]$tls_id_knn)
plot(ldata[[“ToySample”]]$x, ldata[[“ToySample”]]$y,
    col = ifelse(ldata[[“ToySample”]]$tls_id_knn > 0, “red”, “gray”),
    pch = 19, cex = 0.5, main = “Detected TLS (70% sample)”)
# Full dataset with default settings
data(toy_ldata)
ldata <-detect_TLS(“ToySample”, k = 30, ldata = toy_ldata)
table(ldata[[“ToySample”]]$tls_id_knn)
plot(ldata[[“ToySample”]]$x, ldata[[“ToySample”]]$y,
    col = ifelse(ldata[[“ToySample”]]$tls_id_knn > 0, “red”, “gray”),
    pch = 19, cex = 0.5, main = “Detected TLS in toy data”)
~~~

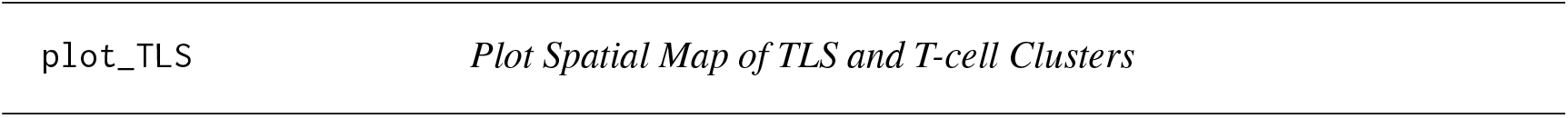

### Description

Produces a ggplot2 scatter plot of cell positions, coloured by TLS membership, T-cell cluster membership, and background phenotype.

Background (non-TLS, non-TIC) cells are rendered with a lower alpha to keep them visually recessive, while TIC cells are drawn slightly larger than TLS cells so they stand out without dominating the plot.

### Usage

~~~
plot_TLS(
   sample,
   ldata = NULL,
   show_tic = TRUE,
   point_size = 0.5,
   alpha = 0.7,
   bg_alpha = 0.25,
   tic_size_mult = 1.8,
   tls_palette = c(“#0072B2”, “#009E73”, “#CC79A7”, “#D55E00”, “#56B4E9”, “#F0E442”),
   tic_colour = “#E69F00”,
   bg_colour = “grey80”
)
~~~

### Arguments

|  |  |
| --- | --- |
| sample | Character. Sample name in ldata. |
| ldata | Named list of data frames, or NULL to use the global ldata object (deprecated; pass explicitly). |
| show_tic | Logical. Colour T-cell clusters (if detect_tic has been run) in a distinct colour? Default TRUE. |
| point_size | Numeric. Base point size for TLS cells and background cells (default 0.5). TIC cells are drawn at point_size * tic_size_mult. |
| alpha | Numeric. Point transparency for TLS and TIC cells (default 0.7). |
| bg_alpha | Numeric. Point transparency for background (non-TLS, non-TIC) cells (default 0.25). Reducing this value pushes background cells further behind the foreground structure. |
| tic_size_mult | Numeric. Multiplier applied to point_size for TIC cells so they appear slightly larger than background and TLS cells (default 1.8). |
| tls_palette | Character vector of colours for TLS IDs. Recycled if there are more TLS than colours. Default uses a colourblind-friendly palette. |
| tic_colour | Character. Colour for T-cell cluster cells (default "#E69F00"). |
| bg_colour | Character. Colour for non-TLS, non-TIC cells (default "grey80"). |

### Value

A `ggplot` object (invisibly). The plot is also printed unless the return value is assigned.

### Examples

~~~
data(toy_ldata)
ldata <-detect_TLS(“ToySample”, k = 30, ldata = toy_ldata)
p <-plot_TLS(“ToySample”, ldata = ldata)
~~~

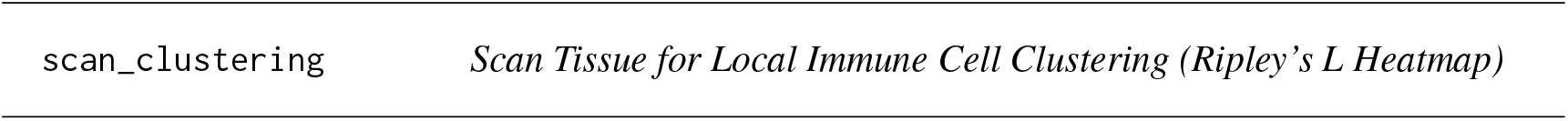

### Description

Applies a sliding-window Ripley’s L analysis across the tissue to produce a spatial clustering map. For each window a *K-integral* index is computed as the mean positive excess of the observed L function over its theoretical CSR value. When plot = TRUE a base-graphics spatial map is drawn with LOESS-smoothed L-excess curves and numeric CI labels overlaid inside each qualifying window, plus a legend identifying point and curve colours.

### Usage

~~~
scan_clustering(
  ws = 500,
  sample,
  phenotype = c(“T cells”, “B cells”, “Both”),
  plot = TRUE,
  creep = 1L,
  min_cells = 10L,
  min_phen_cells = 5L,
  label_cex = 1.1,
  ldata = NULL
)
~~~

### Arguments

|  |  |
| --- | --- |
| `ws` | Numeric. Window side length in microns (default 500). |
| `sample` | Character. Sample name in `ldata`. |
| `phenotype` | One of "T cells", "B cells", or "Both". |
| `plot` | Logical. Draw the spatial clustering map? (default TRUE). |
| `creep` | Integer. Grid density factor; `creep = 2` overlaps adjacent windows by half a window width, producing a smoother map (default 1). |
| `min_cells` | Integer. Minimum total cell count required in a window before it is analysed (default 10). |
| `min_phen_cells` | Integer. Minimum phenotype-specific cell count per window (default 5). |
| `label_cex` | Numeric. Base character expansion for the CI numeric labels drawn inside each window (default 1.1). Increase this value if labels appear too small for your screen or output resolution. |
| `ldata` | Named list of data frames, or NULL to use the global `ldata` object (deprecated; pass explicitly). |

### Details

The K-integral clustering index for window *w* is:

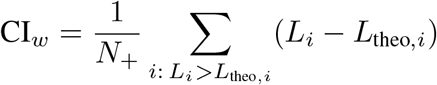

where *N*_+_ is the number of spatial lags where the observed L exceeds the theoretical CSR value. When `plot = TRUE` the map shows:

- All cells as small light-grey points.
- Phenotype cells (T cells green, B cells red).
- Navy dashed grid lines marking window boundaries.
- A LOESS-smoothed L-excess curve inside each qualifying window.
- A bold numeric CI label centred in the window.
- A legend identifying all point and curve colours.

When `phenotype = “Both”` two side-by-side panels are produced - one for B cells and one for T cells - so the two clustering maps can be compared directly on the same spatial layout.

### Value

A named list with elements `B` and/or `T` (depending on phenotype), each containing the Lest objects for all qualifying windows of that phenotype. Returned invisibly when `plot = TRUE`.

### Examples

~~~
data(toy_ldata)
L_models <-scan_clustering(
   ws        = 200,
   sample    = “ToySample”,
   phenotype = “B cells”,
   plot      = TRUE,
   ldata     = toy_ldata
)
cat(“B-cell windows analysed:”, length(L_models$B), “\n”)
# Side-by-side B and T cell panels
L_both <-scan_clustering(
   ws        = 200,
   sample    = “ToySample”,
   phenotype = “Both”,
   plot      = TRUE,
   ldata     = toy_ldata
)
~~~

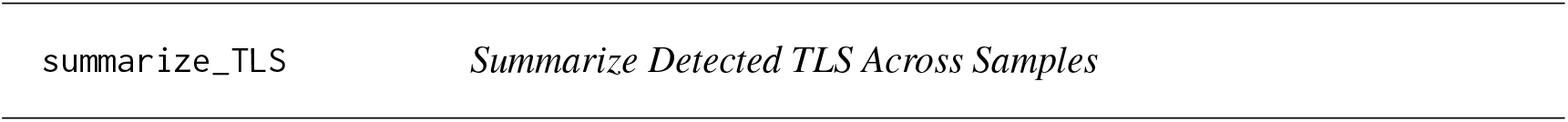

### Description

Produces a tidy `data.frame` with one row per sample summarising the number of detected TLS, their sizes, and (optionally) ICAT scores.

### Usage

summarize_TLS(ldata, calc_icat_scores = FALSE)

### Arguments

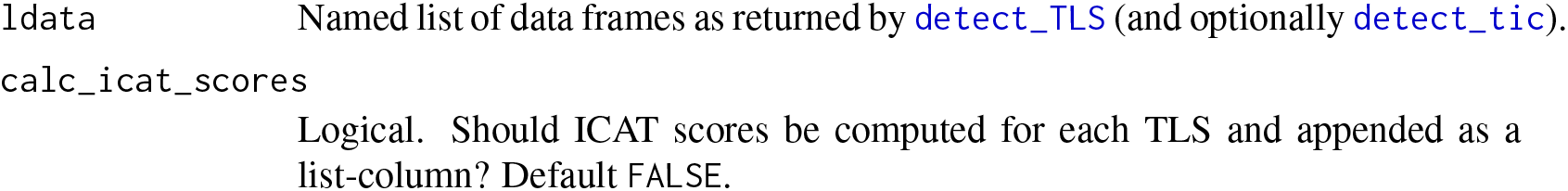

### Value

`A data.frame` with columns:

`sample` Sample name.

`n_TLS` Number of TLS detected.

`total_cells` Total cells in the sample.

`TLS_cells` Number of cells assigned to any TLS.

`TLS_fraction` Fraction of all cells that are TLS cells. mean_TLS_size Mean cells per TLS (`NA` if `n_`TLS = 0).

`n_TIC` Number of T-cell clusters detected by `detect_tic` (NA if not yet run).

`icat_scores` List-column of ICAT scores per TLS (only when `calc_icat_scores = TRUE`).

### Examples

~~~
data(toy_ldata)
ldata <-detect_TLS(“ToySample”, k = 30, ldata = toy_ldata)
summarize_TLS(ldata)
~~~

|  |  |
| --- | --- |
| toy_ldata | *Toy Multiplexed Imaging Data* |

### Description

A small synthetic dataset mimicking multiplexed tissue imaging data, used in package examples and tests. The list contains one sample named “ToySample”.

### Usage

~~~
   toy_ldata
~~~

### Format

A named list with one element:

`ToySample` A `data.frame` with the following columns:

`x` Numeric. X coordinate in microns.

`y` Numeric. Y coordinate in microns.

phenotype Character. Cell phenotype label. Values are `“B cell”, “T cell”`, and `“Other”`.

### Source

Synthetically generated for package examples.

### References

Amiryousefi et al. (2025) doi:10.1101/2025.09.21.677465

### Examples

~~~
data(toy_ldata)
str(toy_ldata)
table(toy_ldata[[“ToySample”]]$phenotype)
~~~

## Index

### *datasets

~~~
  toy_ldata, 11
calc_icat, *2*, 3
detect_tic, *2*, 4, *10*
detect_TLS, *2*, 5, *10*
plot_TLS, *2*, 6
scan_clustering, *2*, 8
summarize_TLS, *2*, 10
tlsR (tlsR-package), 2
tlsR-package, 2
toy_ldata, 11
~~~

## REFERENCES

1. Helmink BA, Reddy SM, Gao J, Zhang S, Basar R, Thakur R, et al. B cells and tertiary lymphoid structures promote immunotherapy response. Nature. 2020 Jan 23;577(7791):549–55. doi:10.1038/s41586-019-1922-8

2. Chen X, Wu P, Liu Z, Li T, Wu J, Zeng Z, et al. Tertiary lymphoid structures and their therapeutic implications in cancer. Cell Oncol Dordr Neth. 2024 Oct;47(5):1579–92. doi:10.1007/s13402-024-00975-1 PubMed PMID: 39133439.

3. Amaria RN, Reddy SM, Tawbi HA, Davies MA, Ross MI, Glitza IC, et al. Neoadjuvant immune checkpoint blockade in high-risk resectable melanoma. Nat Med. 2018 Nov;24(11):1649–54. doi:10.1038/s41591-018-0197-1

4. Hugaboom MB, Wirth LV, Street K, Ruthen N, Jegede OA, Schindler NR, et al. Presence of tertiary lymphoid structures and exhausted tissue-resident T cells determines clinical response to PD-1 blockade in renal cell carcinoma. Cancer Discov. 2025 Feb 24. doi:10.1158/2159-8290.CD-24-0991

5. Chen Y, Wu Y, Yan G, Zhang G. Tertiary lymphoid structures in cancer: maturation and induction. Front Immunol. 2024 Apr 16;15. doi:10.3389/fimmu.2024.1369626

6. Cabrita R, Lauss M, Sanna A, Donia M, Skaarup Larsen M, Mitra S, et al. Tertiary lymphoid structures improve immunotherapy and survival in melanoma. Nature. 2020 Jan 23;577(7791):561–5. doi:10.1038/s41586-019-1914-8

7. Zhang C, Wang XY, Zuo JL, Wang XF, Feng XW, Zhang B, et al. Localization and density of tertiary lymphoid structures associate with molecular subtype and clinical outcome in colorectal cancer liver metastases. J Immunother Cancer. 2023 Feb 9;11(2). doi:10.1136/jitc-2022-006425 PubMed PMID: 10.1136/jitc-2022-006425.

8. Abida W, Cheng ML, Armenia J, Middha S, Autio KA, Vargas HA, et al. Analysis of the Prevalence of Microsatellite Instability in Prostate Cancer and Response to Immune Checkpoint Blockade. JAMA Oncol. 2019 Apr 1;5(4):471. doi:10.1001/jamaoncol.2018.5801

9. Irani J, Goujon JM, Ragni E, Peyrat L, Hubert J, Saint F, et al. High-grade inflammation in prostate cancer as a prognostic factor for biochemical recurrence after radical prostatectomy. Urology. 1999 Sep;54(3):467–72. doi:10.1016/S0090-4295(99)00152-1

10. Vesalainen S, Lipponen P, Talja M, Syrjänen K. Histological grade, perineural infiltration, tumour-infiltrating lymphocytes and apoptosis as determinants of long-term prognosis in prostatic adenocarcinoma. Eur J Cancer. 1994 Jan;30(12):1797–803. doi:10.1016/0959-8049(94)E0159-2

11. Shahait M, Hakansson AK, Daniel RE, Hosny K, Davicioni E, Liu SY, et al. Quantification and molecular correlates of tertiary lymphoid structures in primary prostate cancer. The Prostate. 2024 Jun;84(8):709–16. doi:10.1002/pros.24684

12. McArdle PA, Canna K, McMillan DC, McNicol AM, Campbell R, Underwood MA. The relationship between T-lymphocyte subset infiltration and survival in patients with prostate cancer. Br J Cancer. 2004 Aug;91(3):541–3. doi:10.1038/sj.bjc.6601943

13. Mercader M, Bodner BK, Moser MT, Kwon PS, Park ES, Manecke RG, et al. T cell infiltration of the prostate induced by androgen withdrawal in patients with prostate cancer. Proc Natl Acad Sci U S A. 2001 Dec 4;98(25):14565–70. doi:10.1073/pnas.251140998 PubMed PMID: 11734652; PubMed Central PMCID: PMC64722.

14. Ebelt K, Babaryka G, Figel AM, Pohla H, Buchner A, Stief CG, et al. Dominance of CD4+ lymphocytic infiltrates with disturbed effector cell characteristics in the tumor microenvironment of prostate carcinoma. The Prostate. 2008 Jan;68(1):1–10. doi:10.1002/pros.20661

15. Gannon PO, Poisson AO, Delvoye N, Lapointe R, Mes-Masson AM, Saad F. Characterization of the intra-prostatic immune cell infiltration in androgen-deprived prostate cancer patients. J Immunol Methods. 2009 Aug;348(1–2):9–17. doi:10.1016/j.jim.2009.06.004

16. Flammiger A, Bayer F, Cirugeda‐Kühnert A, Huland H, Tennstedt P, Simon R, et al. Intratumoral T but not B lymphocytes are related to clinical outcome in prostate cancer. APMIS. 2012 Nov;120(11):901–8. doi:10.1111/j.1600-0463.2012.02924.x

17. Ness N, Andersen S, Valkov A, Nordby Y, Donnem T, Al-Saad S, et al. Infiltration of CD8+ lymphocytes is an independent prognostic factor of biochemical failure-free survival in prostate cancer: CD8+ Lymphocytes in Prostate Cancer. The Prostate. 2014 Oct;74(14):1452–61. doi:10.1002/pros.22862

18. Zeigler-Johnson C, Morales KH, Lal P, Feldman M. The Relationship between Obesity, Prostate Tumor Infiltrating Lymphocytes and Macrophages, and Biochemical Failure. Boissonnas A, editor. PLOS ONE. 2016 Aug 3;11(8):e0159109. doi:10.1371/journal.pone.0159109

19. Mo R, Han Z, Liang Y, Ye J, Wu S, Lin SX, et al. Expression of PD‐L1 in tumor‐associated nerves correlates with reduced CD8+ tumor‐associated lymphocytes and poor prognosis in prostate cancer. Int J Cancer. 2019 Jun 15;144(12):3099–110. doi:10.1002/ijc.32061

20. Petitprez F, Fossati N, Vano Y, Freschi M, Becht E, Lucianò R, et al. PD-L1 Expression and CD8+ T-cell Infiltrate are Associated with Clinical Progression in Patients with Node-positive Prostate Cancer. Eur Urol Focus. 2019 Mar;5(2):192–6. doi:10.1016/j.euf.2017.05.013

21. Anton A, Hutchinson R, Hovens CM, Christie M, Ryan A, Gibbs P, et al. An immune suppressive tumor microenvironment in primary prostate cancer promotes tumor immune escape. Nie D, editor. PLOS ONE. 2024 Nov 27;19(11):e0301943. doi:10.1371/journal.pone.0301943

22. Woo JR, Liss MA, Muldong MT, Palazzi K, Strasner A, Ammirante M, et al. Tumor infiltrating B-cells are increased in prostate cancer tissue. J Transl Med. 2014 Dec;12(1):30. doi:10.1186/1479-5876-12-30

23. Valdman A, Jaraj SJ, Compérat E, Charlotte F, Roupret M, Pisa P, et al. Distribution of Foxp3-, CD4-and CD8-positive lymphocytic cells in benign and malignant prostate tissue. APMIS Acta Pathol Microbiol Immunol Scand. 2010 May;118(5):360–5. doi:10.1111/j.1600-0463.2010.02604.x PubMed PMID: 20477811.

24. Vicier C, Ravi P, Kwak L, Werner L, Huang Y, Evan C, et al. Association between CD8 and PD‐L1 expression and outcomes after radical prostatectomy for localized prostate cancer. The Prostate. 2021 Jan;81(1):50–7. doi:10.1002/pros.24079

25. Flammiger A, Weisbach L, Huland H, Tennstedt P, Simon R, Minner S, et al. High tissue density of FOXP3+ T cells is associated with clinical outcome in prostate cancer. Eur J Cancer. 2013 Apr;49(6):1273–9. doi:10.1016/j.ejca.2012.11.035

26. Gollapudi K, Galet C, Grogan T, Zhang H, Said JW, Huang J, et al. Association between tumor-associated macrophage infiltration, high grade prostate cancer, and biochemical recurrence after radical prostatectomy. Am J Cancer Res. 2013;3(5):523–9. PubMed PMID: 24224130; PubMed Central PMCID: PMC3816972.

27. Richardsen E, Uglehus RD, Due J, Busch C, Busund LR. The prognostic impact of M‐CSF, CSF‐1 receptor, CD68 and CD3 in prostatic carcinoma. Histopathology. 2008 Jul;53(1):30–8. doi:10.1111/j.1365-2559.2008.03058.x

28. Andersen LB, Nørgaard M, Rasmussen M, Fredsøe J, Borre M, Ulhøi BP, et al. Immune cell analyses of the tumor microenvironment in prostate cancer highlight infiltrating regulatory T cells and macrophages as adverse prognostic factors. J Pathol. 2021 Oct;255(2):155–65. doi:10.1002/path.5757 PubMed PMID: 34255349.

29. Calagua C, Ficial M, Jansen CS, Hirz T, Del Balzo L, Wilkinson S, et al. A Subset of Localized Prostate Cancer Displays an Immunogenic Phenotype Associated with Losses of Key Tumor Suppressor Genes. Clin Cancer Res. 2021 Sep 1;27(17):4836–47. doi:10.1158/1078-0432.CCR-21-0121

30. García-Hernández MDLL, Uribe-Uribe NO, Espinosa-González R, Kast WM, Khader SA, Rangel-Moreno J. A Unique Cellular and Molecular Microenvironment Is Present in Tertiary Lymphoid Organs of Patients with Spontaneous Prostate Cancer Regression. Front Immunol. 2017 May 17;8:563. doi:10.3389/fimmu.2017.00563

31. Ak Ç, Sayar Z, Thibault G, Burlingame EA, Kuykendall MJ, Eng J, et al. Multiplex imaging of localized prostate tumors reveals altered spatial organization of AR-positive cells in the microenvironment. iScience. 2024 Sep 20;27(9):110668. doi:10.1016/j.isci.2024.110668 PubMed PMID: 39246442; PubMed Central PMCID: PMC11379676.

32. Lin JR, Wang S, Coy S, Chen YA, Yapp C, Tyler M, et al. Multiplexed 3D atlas of state transitions and immune interaction in colorectal cancer. Cell. 2023 Jan;186(2):363–381.e19. doi:10.1016/j.cell.2022.12.028

33. Lin JR, Izar B, Wang S, Yapp C, Mei S, Shah PM, et al. Highly multiplexed immunofluorescence imaging of human tissues and tumors using t-CyCIF and conventional optical microscopes. eLife. 2018 Jul 11;7:e31657. doi:10.7554/eLife.31657

34. Pinsky PF, Miller EA, P p, R G, Ed C G A. Extended follow-up for prostate cancer incidence and mortality in the PLCO randomized cancer screening trial. BJU Int. 2019 May;123(5):854–60. doi:10.1111/bju.14580 PubMed PMID: 30288918; PubMed Central PMCID: PMC6450783.

35. Borregales LD, DeMeo G, Gu X, Cheng E, Dudley V, Schaeffer EM, et al. Grade Migration of Prostate Cancer in the United States During the Last Decade. JNCI J Natl Cancer Inst. 2022 Mar 28;114(7):1012–9. doi:10.1093/jnci/djac066 PubMed PMID: 35348709; PubMed Central PMCID: PMC9275764.

36. Schapiro D, Sokolov A, Yapp C, Chen YA, Muhlich JL, Hess J, et al. MCMICRO: a scalable, modular image-processing pipeline for multiplexed tissue imaging. Nat Methods. 2022 Mar;19(3):3. doi:10.1038/s41592-021-01308-y PubMed PMID: 34824477; PubMed Central PMCID: PMC8916956.

37. Baker GJ, Novikov E, Zhao Z, Vallius T, Davis JA, Lin JR, et al. Quality control for single-cell analysis of high-plex tissue profiles using CyLinter. Nat Methods. 2024 Dec;21(12):2248–59. doi:10.1038/s41592-024-02328-0

38. Vinuesa C, Shen Q, Wang H, Meng X, Roco J, Battaglia M, et al. The transcription factors TCF1 and LEF1 drive B-1a cell development, self-renewal, and regulatory function. [Internet]. Research Square; 2024 [cited 2025 Aug 10]. Available from: https://www.researchsquare.com/article/rs-3855081/v1 doi:10.21203/rs.3.rs-3855081/v1

39. Baker GJ, Novikov E, Coy S, Chen YA, Hug CB, Ahmed Z, et al. Morphology-Aware Profiling of Highly Multiplexed Tissue Images using Variational Autoencoders [Internet]. bioRxiv; 2025 [cited 2026 Jun 2]. p. 2025.06.23.661064. Available from: https://www.biorxiv.org/content/10.1101/2025.06.23.661064v2 doi:10.1101/2025.06.23.661064

40. Kiskowski MA, Hancock JF, Kenworthy AK. On the Use of Ripley’s K-Function and Its Derivatives to Analyze Domain Size. Biophys J. 2009 Aug 19;97(4):1095–103. doi:10.1016/j.bpj.2009.05.039 PubMed PMID: 19686657; PubMed Central PMCID: PMC2726315.

41. Feng Y, Yang T, Zhu J, Li M, Doyle M, Ozcoban V, et al. Spatial analysis with SPIAT and spaSim to characterize and simulate tissue microenvironments. Nat Commun. 2023 May 15;14(1):2697. doi:10.1038/s41467-023-37822-0 PubMed PMID: 37188662; PubMed Central PMCID: PMC10185501.

42. Kermany DS, Ahn JY, Vasquez M, Zhang W, Wang L, Liu K, et al. Multiscale 3D spatial analysis of the tumor microenvironment using whole‐tissue digital histopathology. Cancer Commun. 2025 Jan 3;45(3):386–90. doi:10.1002/cac2.12655 PubMed PMID: 39749685; PubMed Central PMCID: PMC11947609.

43. Hyvärinen A, Oja E. Independent component analysis: algorithms and applications. Neural Netw Off J Int Neural Netw Soc. 2000;13(4–5):411–30. doi:10.1016/s0893-6080(00)00026-5 PubMed PMID: 10946390.

44. Fridman WH, Meylan M, Pupier G, Calvez A, Hernandez I, Sautès-Fridman C. Tertiary lymphoid structures and B cells: An intratumoral immunity cycle. Immunity. 2023 Oct 10;56(10):2254–69. doi:10.1016/j.immuni.2023.08.009

45. Lin JR, Chen YA, Campton D, Cooper J, Coy S, Yapp C, et al. High-plex immunofluorescence imaging and traditional histology of the same tissue section for discovering image-based biomarkers. Nat Cancer. 2023 Jul;4(7):1036–52. doi:10.1038/s43018-023-00576-1

46. Ripley BD. The second-order analysis of stationary point processes. J Appl Probab. 1976 Jun;13(2):255–66. doi:10.2307/3212829

47. Baddeley A, Rubak E, Turner R. Spatial point patterns: methodology and applications with R. Boca Raton ; London ; New York: CRC Press, Taylor & Francis Group; 2016. 810 p. (Champan & Hall/CRC Interdisciplinary Statistics Series).

48. Discussion on Dr Ripley’s Paper. J R Stat Soc Ser B Stat Methodol. 1977 Jan 1;39(2):192–212. doi:10.1111/j.2517-6161.1977.tb01616.x

49. Møller J, Waagepetersen RP. Statistical inference and simulation for spatial point processes. Boca Raton: Chapman & Hall/CRC; 2004. 300 p.

50. Epstein JI, Egevad L, Amin MB, Delahunt B, Srigley JR, Humphrey PA, et al. The 2014 International Society of Urological Pathology (ISUP) Consensus Conference on Gleason Grading of Prostatic Carcinoma: Definition of Grading Patterns and Proposal for a New Grading System. Am J Surg Pathol. 2016 Feb;40(2):244–52. doi:10.1097/PAS.0000000000000530 PubMed PMID: 26492179.

51. Lin JR, Izar B, Wang S, Yapp C, Mei S, Shah PM, et al. Highly multiplexed immunofluorescence imaging of human tissues and tumors using t-CyCIF and conventional optical microscopes. Chakraborty AK, Raj A, Marr C, Horváth P, editors. eLife. 2018 Jul 11;7:e31657. doi:10.7554/eLife.31657

52. Le DT, Uram JN, Wang H, Bartlett BR, Kemberling H, Eyring AD, et al. PD-1 Blockade in Tumors with Mismatch-Repair Deficiency. N Engl J Med. 2015 Jun 25;372(26):2509–20. doi:10.1056/NEJMoa1500596 PubMed PMID: 26028255.

53. Yang S, Yang Y, Fang Y, Zhou Q, Sun W, Zhang Z, et al. Targeting tumour-infiltrating B cells: mechanisms and advances in cancer therapy. Cell Death Dis. 2025 Nov 24;17(1):53. doi:10.1038/s41419-025-08254-z

54. Yang DD, Abdelnaser A, Haas AJ, Wala J, Barney AA, Saad E, et al. Immune Spatial Organization Predicts Distant Metastasis Risk in Aggressive Localized Prostate Cancer. Clin Cancer Res. 2026 May 11. doi:10.1158/1078-0432.CCR-25-4900

55. Nirmal AJ, Maliga Z, Vallius T, Quattrochi B, Chen AA, Jacobson CA, et al. The Spatial Landscape of Progression and Immunoediting in Primary Melanoma at Single-Cell Resolution. Cancer Discov. 2022 Jun 2;12(6):1518–41. doi:10.1158/2159-8290.CD-21-1357

56. Chen Z, Soifer I, Hilton H, Keren L, Jojic V. Modeling Multiplexed Images with Spatial-LDA Reveals Novel Tissue Microenvironments. J Comput Biol J Comput Mol Cell Biol. 2020 Aug;27(8):1204–18. doi:10.1089/cmb.2019.0340 PubMed PMID: 32243203; PubMed Central PMCID: PMC7415889.

57. Dieu-Nosjean MC, Goc J, Giraldo NA, Sautès-Fridman C, Fridman WH. Tertiary lymphoid structures in cancer and beyond. Trends Immunol. 2014 Nov;35(11):571–80. doi:10.1016/j.it.2014.09.006 PubMed PMID: 25443495.

58. Im SJ, Obeng RC, Nasti TH, McManus D, Kamphorst AO, Gunisetty S, et al. Characteristics and anatomic location of PD-1+ TCF1+ stem-like CD8 T cells in chronic viral infection and cancer. Proc Natl Acad Sci. 2023 Oct 10;120(41):e2221985120. doi:10.1073/pnas.2221985120

59. A Density-Based Algorithm for Discovering Clusters in Large Spatial Databases with Noise - ADS [Internet]. [cited 2025 Jul 27]. Available from: https://ui.adsabs.harvard.edu/abs/1996kddm.conf.226E/abstract

60. Kinker GS, Vitiello GAF, Ferreira WAS, Chaves AS, Cordeiro de Lima VC, Medina T da S. B Cell Orchestration of Anti-tumor Immune Responses: A Matter of Cell Localization and Communication. Front Cell Dev Biol. 2021;9:678127. doi:10.3389/fcell.2021.678127 PubMed PMID: 34164398; PubMed Central PMCID: PMC8215448.

61. Zhang E, Ding C, Li S, Zhou X, Aikemu B, Fan X, et al. Roles and mechanisms of tumour-infiltrating B cells in human cancer: a new force in immunotherapy. Biomark Res. 2023 Mar 9;11(1):28. doi:10.1186/s40364-023-00460-1 PubMed PMID: 36890557; PubMed Central PMCID: PMC9997025.

62. Van Meerhaeghe T, Néel A, Brouard S, Degauque N. Regulation of CD8 T cell by B-cells: A narrative review. Front Immunol. 2023 Mar 8;14:1125605. doi:10.3389/fimmu.2023.1125605

63. van Rijthoven M, Obahor S, Pagliarulo F, van den Broek M, Schraml P, Moch H, et al. Multi-resolution deep learning characterizes tertiary lymphoid structures and their prognostic relevance in solid tumors. Commun Med. 2024 Jan 5;4(1):5. doi:10.1038/s43856-023-00421-7 PubMed PMID: 38182879; PubMed Central PMCID: PMC10770129.

64. Dieu-Nosjean MC, editor. Tertiary Lymphoid Structures: Methods and Protocols. 2nd ed. 2025. New York, NY: Springer US; 2025. 1 p. (Methods in Molecular Biology; no. 2864). doi:10.1007/978-1-0716-4184-2

65. Rekoske BT, McNeel DG. Immunotherapy for prostate cancer: False promises or true hope? Cancer. 2016 Dec 1;122(23):3598–607. doi:10.1002/cncr.30250 PubMed PMID: 27649312; PubMed Central PMCID: PMC5115970.

66. Yapp C, Nirmal AJ, Zhou FY, Wong AYH, Tefft J, Lu YD, et al. Highly Multiplexed 3D Profiling of Cell States and Immune Niches in Human Tumours [Internet]. bioRxiv; 2025 [cited 2025 Apr 13]. p. 2023.11.10.566670. Available from: https://www.biorxiv.org/content/10.1101/2023.11.10.566670v4 doi:10.1101/2023.11.10.566670

67. Im SJ, Hashimoto M, Gerner MY, Lee J, Kissick HT, Burger MC, et al. Defining CD8+ T cells that provide the proliferative burst after PD-1 therapy. Nature. 2016 Sep 15;537(7620):417–21. doi:10.1038/nature19330 PubMed PMID: 27501248; PubMed Central PMCID: PMC5297183.

68. Siddiqui I, Schaeuble K, Chennupati V, Fuertes Marraco SA, Calderon-Copete S, Pais Ferreira D, et al. Intratumoral Tcf1+PD-1+CD8+ T Cells with Stem-like Properties Promote Tumor Control in Response to Vaccination and Checkpoint Blockade Immunotherapy. Immunity. 2019 Jan 15;50(1):195–211.e10. doi:10.1016/j.immuni.2018.12.021 PubMed PMID: 30635237.

69. Fridman WH, Meylan M, Petitprez F, Sun CM, Italiano A, Sautès-Fridman C. B cells and tertiary lymphoid structures as determinants of tumour immune contexture and clinical outcome. Nat Rev Clin Oncol. 2022 Jul;19(7):441–57. doi:10.1038/s41571-022-00619-z

70. Helmink BA, Reddy SM, Gao J, Zhang S, Basar R, Thakur R, et al. B cells and tertiary lymphoid structures promote immunotherapy response. Nature. 2020 Jan 23;577(7791):549–55. doi:10.1038/s41586-019-1922-8

71. Schumacher TN, Thommen DS. Tertiary lymphoid structures in cancer. Science. 2022 Jan 7;375(6576):eabf9419. doi:10.1126/science.abf9419 PubMed PMID: 34990248.

72. Vanhersecke L, Bougouin A, Crombé A, Brunet M, Sofeu C, Parrens M, et al. Standardized Pathology Screening of Mature Tertiary Lymphoid Structures in Cancers. Lab Invest. 2023 May 1;103(5):100063. doi:10.1016/j.labinv.2023.100063

73. Wang Y, Lin H, Yao N, Chen X, Qiu B, Cui Y, et al. Computerized tertiary lymphoid structures density on H&E-images is a prognostic biomarker in resectable lung adenocarcinoma. iScience. 2023 Sep 15;26(9). doi:10.1016/j.isci.2023.107635

74. Nirmal AJ, Sorger PK. SCIMAP: A Python Toolkit for Integrated Spatial Analysis of Multiplexed Imaging Data. J Open Source Softw. 2024 May 29;9(97):6604. doi:10.21105/joss.06604

75. Jeong S hwan, Kwak C. Immunotherapy for prostate cancer: Requirements for a successful regime transfer. Investig Clin Urol. 2022 Jan;63(1):3–13. doi:10.4111/icu.20210369 PubMed PMID: 34983117; PubMed Central PMCID: PMC8756154.

76. Beer TM, Kwon ED, Drake CG, Fizazi K, Logothetis C, Gravis G, et al. Randomized, Double-Blind, Phase III Trial of Ipilimumab Versus Placebo in Asymptomatic or Minimally Symptomatic Patients With Metastatic Chemotherapy-Naive Castration-Resistant Prostate Cancer. J Clin Oncol Off J Am Soc Clin Oncol. 2017 Jan;35(1):40–7. doi:10.1200/JCO.2016.69.1584 PubMed PMID: 28034081.

77. Ozbek B, Ertunc O, Erickson A, Vidal ID, Gomes‐Alexandre C, Guner G, et al. Multiplex immunohistochemical phenotyping of T cells in primary prostate cancer. The Prostate. 2022 May;82(6):706–22. doi:10.1002/pros.24315

78. Wu Z, Chen H, Luo W, Zhang H, Li G, Zeng F, et al. The Landscape of Immune Cells Infiltrating in Prostate Cancer. Front Oncol. 2020 Oct 29;10:517637. doi:10.3389/fonc.2020.517637

79. Hirz T, Mei S, Sarkar H, Kfoury Y, Wu S, Verhoeven BM, et al. Dissecting the immune suppressive human prostate tumor microenvironment via integrated single-cell and spatial transcriptomic analyses. Nat Commun. 2023 Feb 7;14(1):663. doi:10.1038/s41467-023-36325-2

80. Song H, Weinstein HNW, Allegakoen P, Wadsworth MH, Xie J, Yang H, et al. Single-cell analysis of human primary prostate cancer reveals the heterogeneity of tumor-associated epithelial cell states. Nat Commun. 2022 Jan 10;13(1):141. doi:10.1038/s41467-021-27322-4

81. Rebuzzi SE, Rescigno P, Catalano F, Mollica V, Vogl UM, Marandino L, et al. Immune Checkpoint Inhibitors in Advanced Prostate Cancer: Current Data and Future Perspectives. Cancers. 2022 Feb 28;14(5):1245. doi:10.3390/cancers14051245 PubMed PMID: 35267553; PubMed Central PMCID: PMC8909751.

82. Leone G, Wong YNS, Jones RJ, Sankey P, Josephs DH, Crabb SJ, et al. Nivolumab and Ipilimumab for Metastatic Castration-Resistant Prostate Cancer With an Immunogenic Signature: The Multicenter, Two-Cohort, Phase II NEPTUNES Study. J Clin Oncol Off J Am Soc Clin Oncol. 2025 Oct;43(28):3070–80. doi:10.1200/JCO-24-02637 PubMed PMID: 40720716.

